# Learning human-robot collaboration reshapes biological motor coordination

**DOI:** 10.64898/2026.11.11.765896

**Authors:** Ema Jugovic, Peiqi Kang, Julien Russ, Celia Foster, Viktorija Pavalkyte, Zehra Merchant, Martina Giancane, Rupsha Panda, Eden Lechelle-Rambach, Juan Alvaro Gallego, Hristo Dimitrov, Gauthier Everard, Dani Clode, Tamar R. Makin

**Affiliations:** Medical Research Council Cognition and Brain Sciences Unit, University of Cambridge, Cambridge, UK; The BioRobotics Institute and Department of Excellence in Robotics and AI, Scuola Superiore Sant’Anna, Pisa, Italy; Department of Psychology, University of Michigan, Ann Arbor, MI, USA; Champalimaud Foundation, Lisbon, Portugal; NeuroMusculoSkeletal Lab, Institut de Recherche Expérimentale et Clinique, UCLouvain, Brussels, Belgium

## Abstract

Wearable augmentation robots expand human motor capabilities, but every added degree of freedom creates a new coordination problem for the biological body. Although previous studies have made substantial advances in robot sensing, control, and safety, they have largely overlooked how the user’s own movement changes: individuals adapt their body movements when they start using an augmentation robot, even when control does not require it, and this coordination pattern continues to change as they gain proficiency. Consequently, users may struggle to accommodate robot-related restructuring of their movements during prolonged use, potentially increasing the risk of user rejection or abandonment. Over five days of training, 26 participants performed tasks with a wearable robotic digit and a tool-use control task, allowing us to longitudinally characterise and compare body-wide movement patterns between the two. Performance improved similarly across tasks, but the underlying motor coordination trajectories diverged. During augmentation, biological finger and limb coordination departed progressively further from its initial structure than during tool use, and joints increasingly varied together in ways that kept task performance stable, indicating that a new, body-wide augmented motor pattern continued to develop across days. These findings demonstrate that the biological motor system is not a fixed substrate that can be simply augmented with artificial limbs, but rather its movement patterns continuously evolve as users learn to control an added effector. Characterising this evolving motor solution informs the design, training, and evaluation of augmentation systems, supporting the sustained integration of added effectors into the human motor system.

**One-Sentence Summary:** Robots that work with the body face a moving target, as collaborating with one teaches people entirely new ways to move.

## Introduction

From Leonardo da Vinci’s designs for winged flying machines and early conceptualisations of spring-driven prostheses to today’s supernumerary robotic arms and supplementary digits, the ambition to extend human motor function has evolved from imaginative speculation into technological reality (*1–6*). Unlike traditional hand-held tools that replace the functionality of an existing limb, a supernumerary robotic limb (SRL) introduces additional degrees of freedom (DoF) that must be controlled alongside those of the biological body. The motor system must therefore learn to control an artificial effector without compromising the biological actions required for the task (*2*). To avoid disrupting biological movement, SRLs are typically controlled through signals that the task does not otherwise use, either movements of an uninvolved body part, such as the toes, or patterns of joint configuration or muscle activity that leave task-relevant biological movement unchanged, an approach known as null-space control (*2*, *7–10*). Yet sustained interaction with SRLs has rarely been studied beyond short laboratory exposure (*7–11*). Existing evidence shows that novice users can achieve control over one or more additional DoF, but not how the biological motor system changes as augmentation performance matures over extended practice periods. The central unresolved question thus is whether sustained learning preserves or progressively restructures the body-wide coordination present during initial use. Without an answer, both theories of augmentation learning and interfaces calibrated for prolonged use rest on the untested assumption that the initial biological motor solution will endure the test of time.

Refining coordination across biological DoF is itself an objective of motor learning. The motor system exploits redundancy across joints and muscles by organising their contributions into task-dependent synergies: patterns of covariation among muscles or joints (*12*, *13*) that stabilise task-relevant variables (*14*, *15*). Practice can restructure these synergies as a skill develops (*12*, *16–18*). Tool-use provides related evidence for experience-dependent motor learning, because it can alter subsequent arm kinematics (*19–22*). Augmentation likely places a deeper demand on this process because the artificial effector must be coordinated alongside, rather than in place of the biological body, which retains its full functional capacity. Indeed, our previous studies with a two-degree-of-freedom wearable robotic augmentation digit showed that training changes coordination among the biological fingers, alters the neural representation of the biological hand, and supports the formation of a sensory representation of the artificial digit (*23*, *24*). These findings demonstrate that augmentation can alter both biological coordination and brain representations of body parts. While this prior research establishes the capacity for brain plasticity, it does not provide the mechanism through which changes related to artificial limb usage unfold: recalibration of an existing one or the formation of a new control policy?

Motor adaptation recalibrates an established policy in response to altered dynamics or sensorimotor mappings, whereas the slower process of de novo learning develops a new control policy when recalibration cannot generate the required commands (*25–28*). Under adaptation, changes in performance and coordination are largest during early exposure; performance then stabilises, and movements remain largely within the initial coordination structure. De novo learning instead involves continued exploration beyond the existing motor repertoire, progressive restructuring of task-related synergies, and shifts in control strategy across days (*17*, *18*, *28*, *29*). Although both processes may be evoked as individuals repeatedly train with a SRL, their longitudinal signatures can indicate which account better characterises prolonged usage of an augmentation robot: a rapid change that stabilises early on, or does repeated practice continuously restructure a solution that develops across days?

These trajectories matter for how augmentation devices are designed. Most SRL interfaces map a fixed control signal onto device commands, calibrated during initial use, on the assumption that the biological body does not otherwise use this signal. Whether this assumption holds over extended practice is unknown. If users adapt, any change in coordination that draws the control signal into biological movement should be confined to early exposure and settle as performance stabilises. If users instead learn a new control policy, coordination continues to restructure across days, and a signal that was available at calibration may later be recruited for biological movement, producing unintended device commands or constraining natural movement. Null-space interfaces are particularly exposed, because a signal’s independence from task-relevant movement is defined by the coordination present at calibration. Characterising how biological coordination changes over prolonged use is therefore needed both to understand augmentation learning and to design interfaces that remain reliable over time.

Here, we trained participants to use a toe-controlled wearable robotic digit for five consecutive days across a series of augmentation tasks. As control, we compared practice with this SRL to performance of a tool-use control task that was designed to be equally novel: the same participants used a conventional tool in an unfamiliar grip configuration, matching the core motor demands of augmentation while removing the robotic control interface. Using hand, body and SRL movement tracking, we aimed to address three questions. First, as performance improves across five days, does coordination among the biological fingers, wrist and arm remain anchored to the pattern present during initial use, or does it progressively restructure? Across training days, we tracked the structure of inter-joint movement patterns, their speed–accuracy trade-off, and whether this joint variability was organised to keep the task outcome stable. Second, are any longitudinal changes specific to augmentation rather than general practice with a novel effector? We address this by comparing practice with the wearable robotic digit with a demand-matched, equally unfamiliar tool-use control task and by testing whether similar changes emerge when the robotic digit acts without interacting with the biological fingers. Third, does the learnt augmentation solution remain stable when postural or cognitive demands change? We addressed this by having participants perform the augmentation task while standing and while solving arithmetic problems throughout training. Together, these questions address how biological coordination evolves during augmentation training. We find that using an augmentation robot does not simply recalibrate existing movement but progressively restructures coordination across the body, pointing towards augmentation as the learning of a new motor skill.

## Results

### Overview

We designed a 5-day study to determine how the human motor system changes when it learns to control a two-DoF wearable robotic digit, each operated by pressure from one big toe (the Third Thumb, Dani Clode Design; Fig. 1A). The experimental design is presented in Fig. 1B,C. Participants had no previous experience with this device, and the tasks were designed to be sufficiently demanding to prevent ceiling effects and permit measurable improvement across the five training days (Supplementary Vid. 1). In the primary Interaction task, participants used the robotic digit together with a biological finger to grasp and transport a small, slippery cylindrical object from a starting position to designated placeholders (Fig. 1C, Supplementary Vid. 2) (*30*). Alongside this primary task, participants trained on four control tasks. (1) In the demand-matched Tool control task, participants used a conventional hand-held tool to grasp and transfer a slippery flat object into placeholders (Fig. 1C, Supplementary Vid. 3). Critically, participants held the tool in a non-standard grip they had no prior experience with. This preserved the core motor demands of the primary task: a novel precision grip that coordinated a biological finger with an external effector to transport a small object at a self-selected pace. The key difference was the absence of the robotic control interface. (2) In the Independent task (Figure 1C, Supplementary Vid. 4), the toe-controlled robotic digit was used to grasp and place an object without assistance from the biological fingers (*31*, *32*). (3) The Standing task replicated the primary task while standing, altering postural and balance demands (*33*). (4) The Cognitive task added concurrent mental arithmetic to the primary task, which increased cognitive demands (*34*, *35*).

**Figure 1.**
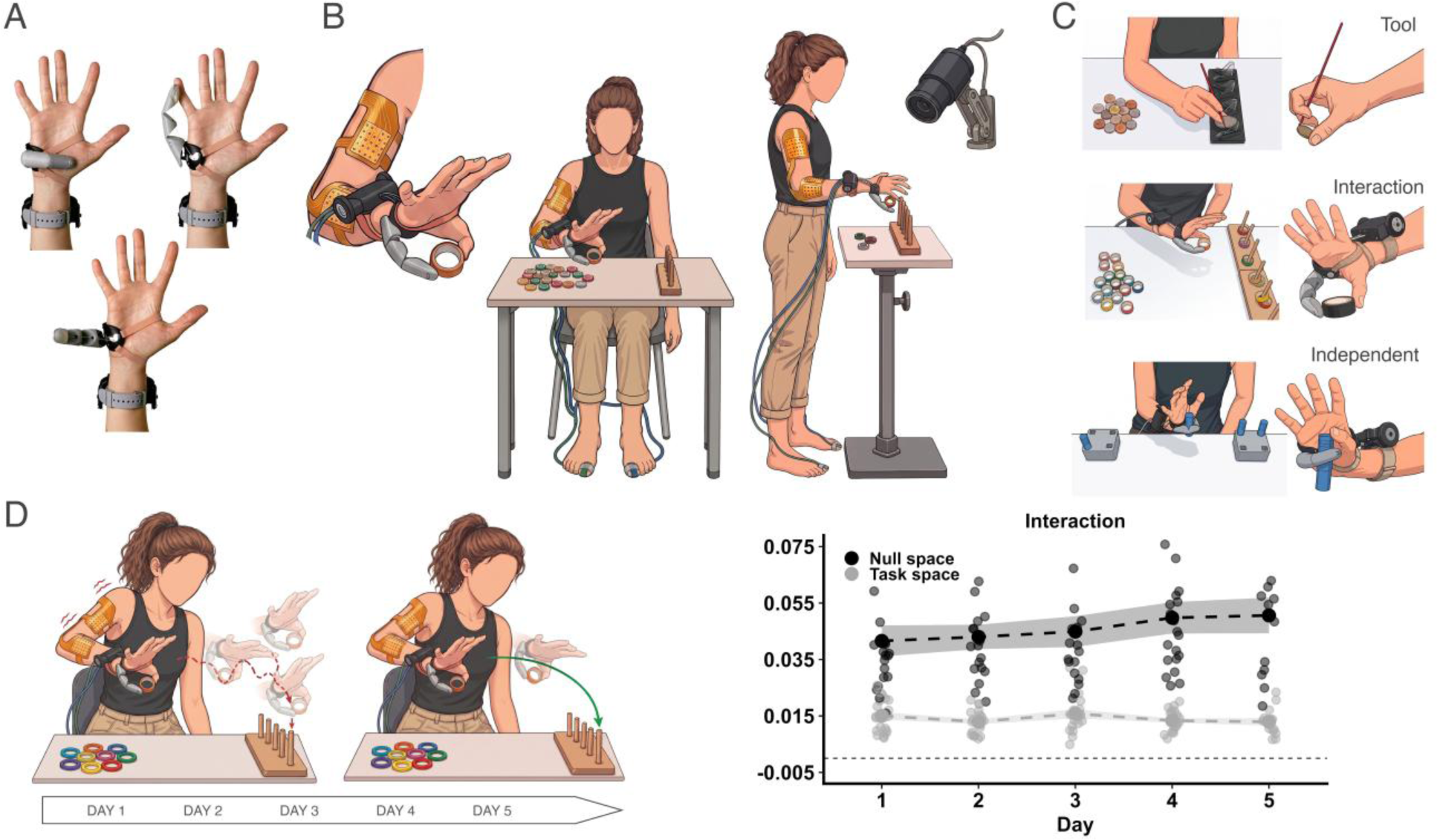
Study design and changes in upper-limb coordination during augmentation training. **A**, The Third Thumb (Dani Clode Design), a 3D-printed robotic digit worn on the ulnar side of the right hand, shown in three configurations across its two (simultaneously used) degrees of freedom. Flexion–extension is controlled by a pressure sensor under the right big toe, and adduction–abduction by a sensor under the left big toe. **B,** Participants trained with the robotic digit while seated and while standing at a height-adjusted table. Upper-limb and hand kinematics were recorded with a markerless multi-camera motion-capture system (FLIR Blackfly S), and muscle activity with a 64-channel custom-made high-density EMG sleeve over the forearm and upper arm. This schematic is for illustration only; see Supplementary materials for full setup details. **C,** Experimental tasks, top to bottom, each shown at the workspace (left) and in close-up (right). Tool: participants picked up coins with a hand-held tool in an unfamiliar grip and placed them into a holder. Interaction: participants grasped tape rolls between the robotic digit and a biological finger and placed them onto wooden poles; this task was also performed while standing and during concurrent mental arithmetics (Cognitive; on Days 1, 3 and 5 only). Independent: participants used the robotic digit alone to extract pegs from one holder and insert them into another. **D,** Schematic summary of changes across five days of Interaction training. Object transport became smoother, and movement exploration decreased over time. Empirical data: null-space variance (*VUCM*; joint combinations that preserve thumb-tip position) increased across training while task-space variance (*VORT*) remained stable, strengthening task-stabilising synergies (detailed below). Dots: individual participants; Large points and dashed lines: group means; Shaded regions: standard error of the mean (SEM); Horizontal dashed line: zero.

Across these tasks, we examined whether improved task performance was accompanied by progressive changes in body-wide kinematics through extended practice. We first quantified improvements in task performance across training, then examined movement smoothness and potential changes in the speed– accuracy relationship.

A total of 26 participants completed five days of training. Markerless motion capture recorded the kinematics of the upper limb, hand, and wearable robot, while a custom-made 64-channel monopolar high-density electromyography (EMG) sleeve recorded muscle activity. After technical exclusions, markerless kinematic data were available for 21 participants. All movement analyses focused on the grasp phase as the main action component of each task. Unless stated otherwise, repeated trial- and block-level observations were aggregated within participants before group summaries were calculated.

### Comparable task performance in augmentation and tool-use masks distinct changes in movement smoothness and speed-accuracy relationships

We compared the Interaction augmentation with the Tool control task to examine whether comparable behavioural learning was accompanied by similar changes in movement control. To establish learning within each task, we min–max normalised daily performance scores within each participant (Fig. 2A). Performance increased across five training days in both the Interaction (*F*(4, 101>) = 42.17, *p* < .0001; Day 1 vs. 5: *t*(82.5) = −11.10, *p* < .0001) and Tool tasks (*F*(4, 102) = 16.69, *p* < .001; Day 1 vs. 5: *t*(82.7) = −7.37, *p* < .0001).

**Figure 2.**
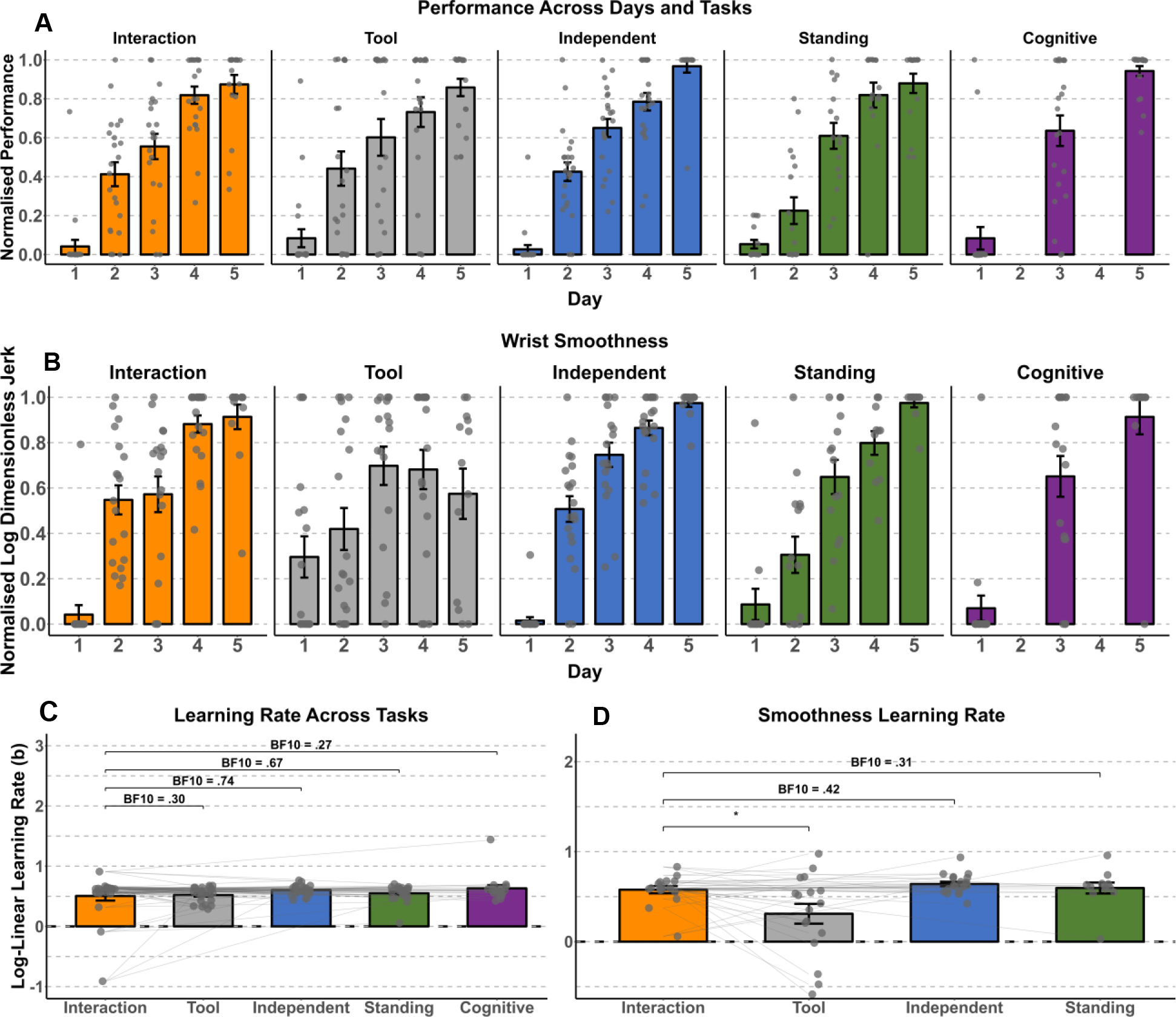
Comparable task learning is accompanied by divergent smoothness trajectories in augmentation and tool-use. **A**, Min–max normalised task performance (0 = worst, 1 = best; within participant and task) increased across all five tasks. **B,** Normalised wrist movement smoothness increased across augmentation tasks, but not in tool-use. **C,** Log-linear task learning rates did not differ between Interaction and any other task (all BF₁₀ < 1, moderately favouring the null). **D,** Log-linear smoothness learning rates likewise differed between the Interaction and Tool tasks. Grey dots and connecting lines: individual participants; coloured bars: group means; error bars: SEM.

Next, we verified that the Interaction and Tool tasks produced comparable gains in task performance by fitting participant-specific logarithmic learning curves to daily scores, min–max normalised within each participant and task to accommodate for their different outcome measures (Fig. 2C). Performance improved similarly in both tasks (Task effect: *F*(1,181.50) = .01, *p* = .939; Day × Task interaction: *F*(4,181.49) = .40, *p* = .812), and the learning slopes provided moderate Bayesian evidence for comparable rates of improvement (mean slopes: Interaction = .55, Tool = .52; paired *t*(21) = .82, *p* = .423; BF₁₀ = .30). Together these results support the comparability of the two tasks in their learning trajectories, providing a common behavioural reference for examining how movement control changed with practice.

We next examined whether comparable behavioural learning was accompanied by similar refinement of the underlying movements. We focused on wrist trajectories as a common kinematic reference for hand transport across tasks. Movement smoothness, a well-established indicator of motor skill, captures the temporal continuity of this transport movement, allowing us to assess whether behavioural improvement coincided with more fluent movement execution. We quantified smoothness using normalised log dimensionless jerk (LDLJ), whose dimensionless formulation accounts for simple differences in movement scale and duration, with higher values indicating smoother movements (Fig. 2B) (*36*, *37*). Wrist smoothness followed distinct training trajectories in the two tasks (Day × Task interaction: *F*(4,168) = 4.64, *p* = .001), with smoothness increasing from Day 1 to 5 in augmentation (paired *t*(158) = −7.43, *p* < 0001) but not in tool-use (paired *t*(159) = −2.35, *p* = .136; Fig. 2B). Tool-use produced smoother wrist movements on Day 1 (paired *t*(151) = 2.38, *p* = .019), but augmentation surpassed it by Day 5 (paired *t*(148) = −2.66, *p* = .009), indicating notable improvement in movement execution only when using the robotic digit, but not when using the tool. Fitted logarithmic learning curves suggested different smoothness acquisition rates (paired *t*(14) = 2.44, *p* = .028; Fig. 2D), providing evidence that augmentation and tool-use refined movement execution at different rates. Despite this kinematic divergence, the two tasks recruited comparable muscle activation patterns. Within-participant cosine similarity between 64-channel MVC-normalised RMS maps from the forearm flexors, forearm extensors, biceps brachii, triceps brachii lateral head, and triceps brachii long head averaged .93 on Day 1 (95% CI: .88 − .98, *n* = 15), well above the conventional .90 threshold for high spatial similarity (*38*, *39*), and remained stable across training days (*F*(4,45.54) = .68, *p* = .610). Comparable behavioural learning and muscular activation thus coexisted with divergent smoothness trajectories, suggesting that the two tasks differed in fine-grained movement refinement rather than in overall motor activity patterns.

The increase in wrist smoothness with practice in the Interaction task raised the question of whether practice also reduced the loss of accuracy that comes with moving faster. We examined this possibility using participant-day slopes relating trajectory error to peak wrist speed (Fig. 3A,B). Speed-accuracy slopes were steeper in the Interaction than the Tool task on Day 1 (paired *t*(129) = −2.20, *p* = .030) but not on Day 5 (paired *t*(123) = −1.04, *p* = .302; Day × Task: *F*(4,123.68) = 2.43, *p* = .051). In the Interaction task, faster movements were significantly associated with greater trajectory error on Day 1 (slope against zero: *t*(64.6) = 3.16, *p* = .002), but not on Day 5 (slope against zero*: t*(71.5) = 1.43, *p* = .158). Tool-use, by contrast, showed no significant speed–accuracy trade-off at either time point (Day 1: *t*(73.7) = .30, *p* = .767, Day 5: *t*(75.2) = .11, *p* = .915). Together, these results suggest that the speed–accuracy trade-off observed early in Interaction training weakened with practice, while task performance continued to improve. One possible interpretation is that the two tasks reflect different learning mechanisms. In adaptation, an existing motor policy is refined without novel skill acquisition, thus no shifts in the speed–accuracy trade-off relationship are expected early in learning, as in the Tool task (*26*, *40*). In de novo learning, a new skill is acquired, and early movements become less accurate as speed increases, while with practice, accuracy at faster speeds improves (*26*, *40*). The early speed–accuracy trade-off in the Interaction task, which was no longer detectable by Day 5, is therefore more indicative of de novo learning, as we elaborate in the Discussion. Augmentation and tool use therefore diverged in movement refinement despite comparable behavioural learning and muscular recruitment. Only in the Interaction task did wrist smoothness increase with practice, and only there did the speed–accuracy trade-off weaken. This combination points towards the formation of a new control policy rather than the adjustment of an existing one, and suggests that augmentation alters the fine-grained temporal structure of movement in ways that tool use does not.

**Figure 3.**
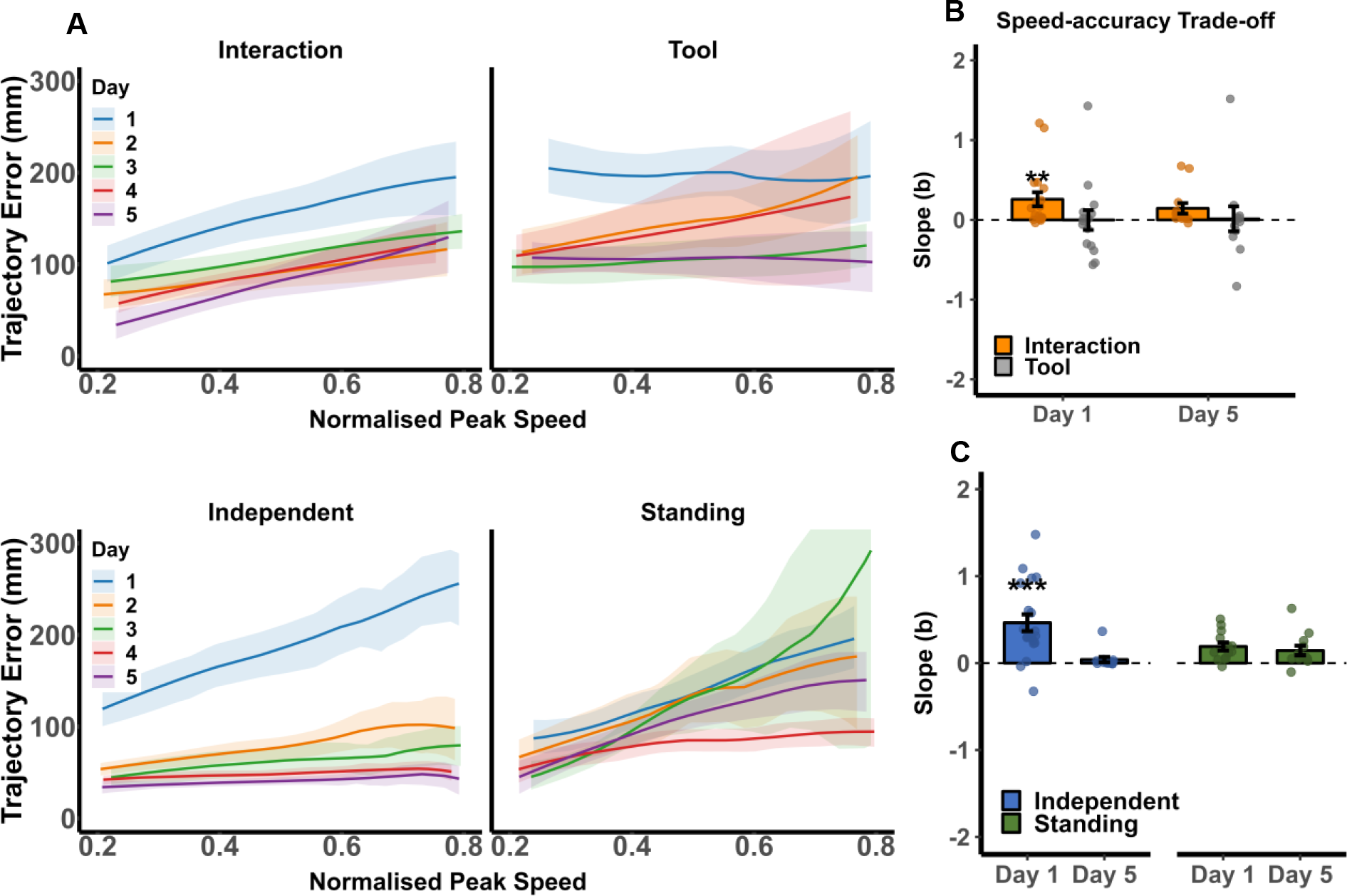
A speed–accuracy trade-off is specific to augmentation despite comparable task learning. **A**, Group-level speed– accuracy curves for the Interaction, Tool (upper panels), Independent and Standing tasks (bottom panels) across Days 1–5. Each curve plots normalised peak wrist speed against trajectory error, defined as the excess wrist path length beyond the shortest task-appropriate path. On Day 1, error increased with speed in the Interaction and Independent tasks, and this dependence weakened with training. **B,** Speed–error slopes (*b*) for the Interaction and Tool tasks on Days 1 and 5, tested against zero. Positive slopes indicate that faster movements produced larger errors. On Day 1, the slope differed significantly from zero in the Interaction task but not in the Tool task. **C,** As in (B), for the Independent and Standing tasks. On Day 1, the slope differed significantly from zero in the Independent task but not in the Standing task. Coloured lines and bars: Group means; Error bars and shaded regions: SEM; ** *p* < .01; *** *p* < .001.

### Distinct restructuring of movement coordination in augmentation and tool-use throughout learning

The proposed distinction between acquiring a new control strategy and refining an existing one led us to examine how finger and arm coordination changed in the Interaction and Tool tasks. We first assessed biological-finger coordination using a principal component analysis (PCA) fitted separately to each participant’s Day 1 data in each task to measure how much variance throughout training remained within the initial coordination patterns. Fisher z-transformed Pearson correlations between metacarpophalangeal (MCP) joint-angle time courses complemented this measure by capturing co-movement between individual fingers (Fig. 4A,B).

**Figure 4.**
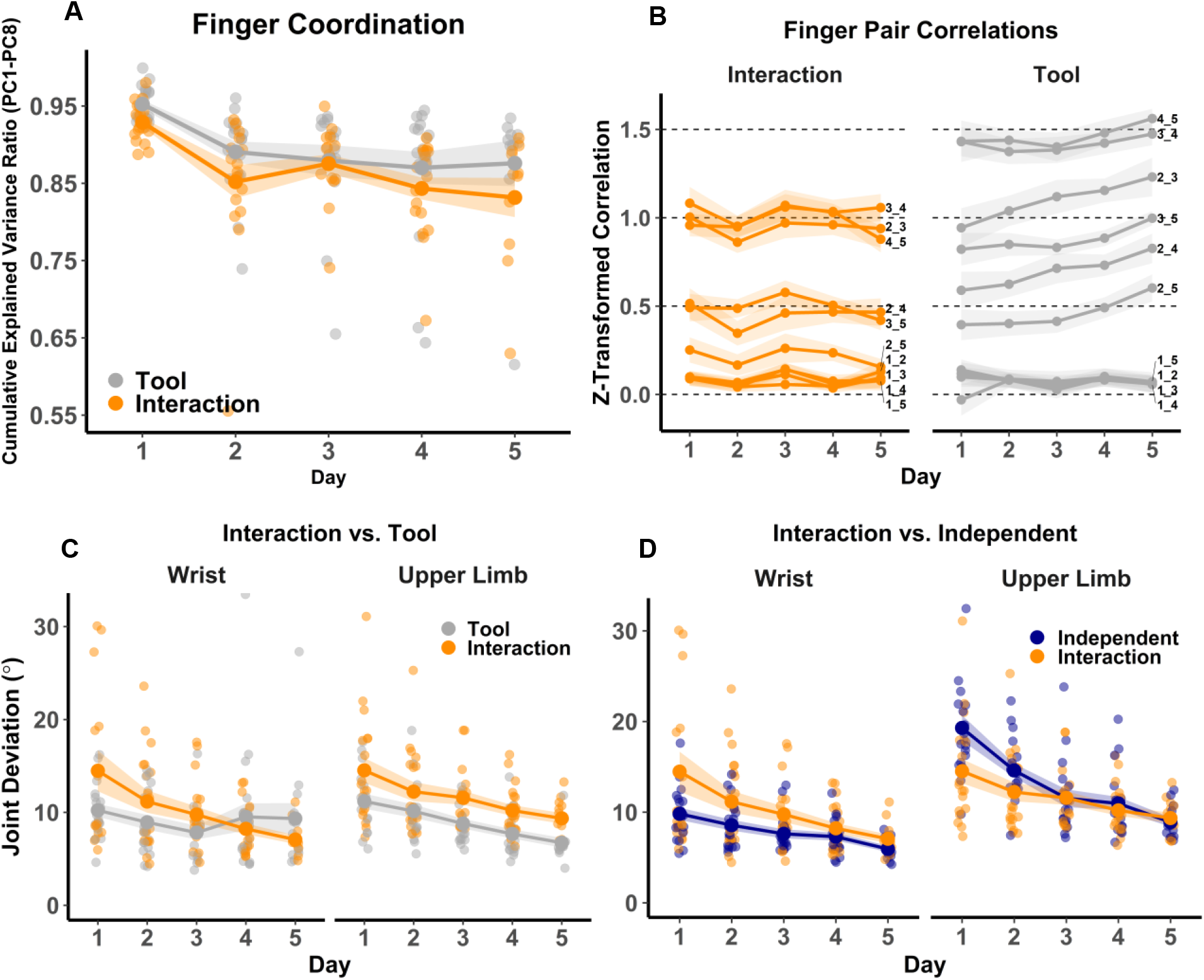
Augmentation loosens finger coupling and drives upper-limb exploration regardless of biological finger involvement. **A**, Cumulative explained variance of the first eight Day 1 PCA components across training. Augmentation drove greater departure from initial finger coordination patterns than tool-use. **B,** Pairwise finger correlations. Augmentation loosened inter-finger coupling, whereas tool-use strengthened it. **C, D,** Joint deviation from each participant’s final-block movement pattern on Day 5 in the Interaction vs. Tool (C) and Interaction vs. Independent (D) tasks. Larger values indicate greater exploration away from the eventual solution. **C,** Augmentation produced greater exploration than tool-use across the upper limb. **D,** Upper-limb exploration exceeded wrist exploration in Independent augmentation. In the first two days, wrist exploration was higher in Interaction than Independent, while upper-limb exploration showed the reverse; both differences resolved by Day 5. Finger labels: 1, biological thumb; 2, index; 3, middle; 4, ring; 5, little finger. Other figure annotations are as detailed above.

The cumulative fraction of variance explained by the first eight Day 1 components decreased across training (Day: *F*(4,136.47) = 10.50, *p* < .0001) and was lower overall in the Interaction task (Task: *F*(1,143.56) = 37.08, *p* = .009). Finger coordination therefore departed from the Day 1 pattern with practice, and lay further from it during Interaction than during tool-use. The greater overall departure in the Interaction task suggests that augmentation reshaped finger coordination more so than tool use did.

Inter-finger correlations told a complementary story: correlations were lower when participants trained on the augmentation device relative to tool-use (Finger Pair × Task interaction: *F*(9,1659.8) = 23.68, *p* < .0001), driven by weaker coupling among the index, middle, and ring fingers, and between these fingers and the pinky (all comparisons *p* < .0001).

The two tasks also diverged over time (Day × Task interaction: *F*(4,1661.3) = 4.75, *p* = .001). In the Tool task, inter-finger correlations increased from Day 1 to Day 5 (paired *t*(1673) = −.3.45, *p* = .001) but decreased from Day 1 to Day 2 in the Interaction task (paired *t*(1665) = 2.02, *p* = .044), and remained stable thereafter. This early reduction in inter-finger correlation, followed by stability, resembles recalibration of an existing inter-finger coupling more than progressive restructuring. Tool use thus strengthened existing biological finger coupling with training, whereas augmentation loosened it early in training before reaching a stable level.

Because changes in hand movement can be supported by wrist and proximal arm motion, we next examined how far these differences extended along the limb. Joint deviation quantified how far each grasp trajectory lay from the participant’s final-block movement pattern on Day 5, with larger values indicating greater exploration away from the final solution (Fig. 4C). Exploration decreased across training (*F*(4,315.69) = 14.85, *p* < .0001) and was greater in augmentation than tool-use (*F*(1,313.19) = 14.50, *p* < .001), particularly at the elbow and shoulder (Task × Body part interaction: (*F*(1,310.34) = 4.04, *p* = .045). In augmentation, upper-limb exploration exceeded wrist exploration (paired *t*(311) = −2.41, *p* = .017), whereas no such difference emerged in tool-use (paired *t*(311) = .44, *p* = .657). Augmentation thus drove broader motor exploration that propagated beyond the device’s point of attachment to the upper limb, suggesting that even a small hand-mounted device can restructure coordination across the entire arm.

Because multiple joint configurations can produce the same end-effector position, not all joint-level variability affects task performance equally. For instance, the same position of the hand or robotic digit can be potentially achieved with different proximal arm configurations. We used uncontrolled manifold (UCM) analysis to decompose angular variance of the biological thumb, wrist, elbow, and shoulder into a null-space, which captured movement variance that did not affect the thumb-tip position (*V_UCM_*) and a task-space component, which captured movements that did affect it (*V_ORT_*). Augmentation and tool-use showed distinct variance redistribution patterns (Fig. 5A). In the Tool task, task-space variance decreased across training (Day × Task interaction: *F*(4, 145.73) = 2.89, *p* = .024; Day 1 vs. 5: paired *t*(150) = 3.56, *p* = .005), while null-space variance remained stable (Day × Task interaction: *F*(4, 139.88) = 2.71; paired *t*(143) = .55, *p* = .982). Augmentation exhibited the opposite pattern where task-space variance remained stable (paired *t*(148) = 1.09, *p* = .811), while null-space variability increased from first to last training day (paired *t*(141) = −3.34, *p* = .009). Although task-space was comparable between tasks on Day 1 (paired *t*(145) = −.58, *p* = .560), it was consistently higher in augmentation than tool-use on all subsequent days (all *p* < .05), while null-space variance was greater in augmentation throughout (all *p* < .05). Augmentation thus did not transform null-space flexibility into task-space precision as tool-use did. This pattern is consistent with an early stage of de novo learning in which participants progressively expanded null-space exploration around the added effector before they start to produce more consistent movements in the task-space (*41*, *42*).

**Figure 5.**
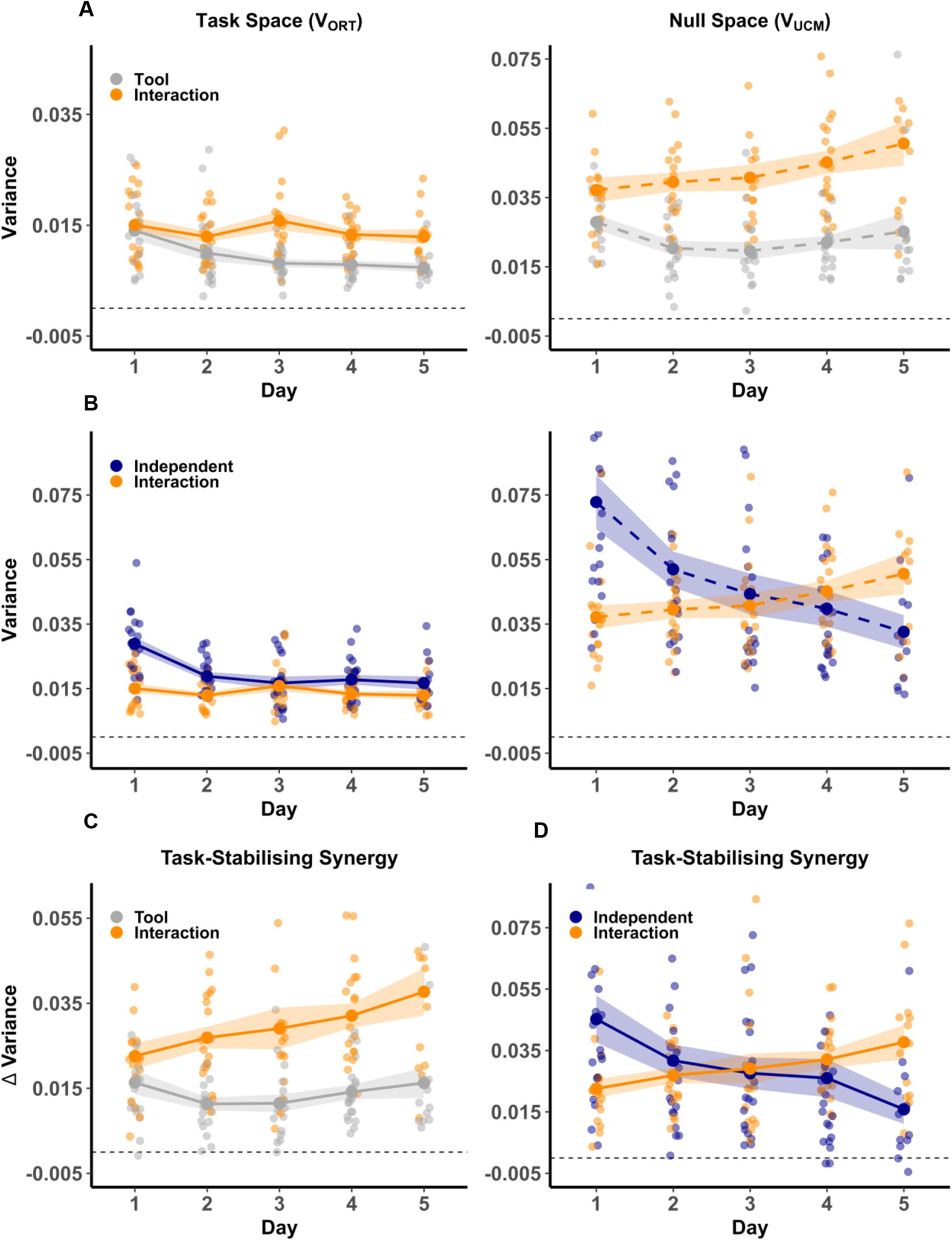
Augmentation selectively expands null-space variance and strengthens synergies, while independent use compresses both subspaces. Joint-angle variance decomposed into task-space (*V*_ORT_, left) and null-space (*V*_UCM_, right) components. A, *V*_ORT_ decreased over training in tool-use but remained stable in augmentation. *V*_UCM_ increased from Day 1 to 5 only in augmentation. **B,** Both *V*_ORT_ and *V*_UCM_ decreased across training in the Independent task, whereas *V*_UCM_ increased in the Interaction task. **C,** Both the Interaction and Tool tasks exhibited task-stabilising synergy (*ΔV* = *V*_UCM_ −*V*_ORT_) from Day 1, but only the Interaction task strengthened it across training. **D,** Task-stabilising synergy increased across training in the Interaction task but decreased in the Independent task, indicating that coordination between biological and robotic digits drives synergy restructuring. Horizontal dashed line: zero. Other figure annotations are as detailed above.

Although augmentation maintained higher absolute task-space variance, joint variability may nonetheless be preferentially structured to stabilise end-effector position. We assessed this using the task-stabilising synergy index *ΔV* = *V_UCM_* -*V_ORT_* , where positive values indicate increased variability in the null-space when compared to the task-space (Fig. 5C). *ΔV* increased from Day 1 to 5 in augmentation (Day × Task interaction: *F*(4, 141.52) = 2.48, *p* = .047; paired *t*(144) = −3.78, *p* = .002) but not in tool-use (paired *t*(145) = .05, *p* = 1.00), and was greater in augmentation on all days except Day 1 (all *p* < .0001). Thus, while augmentation entailed greater overall task- and null-space variability, this variability became increasingly structured to preserve end-effector accuracy across training.

Despite comparable performance gains across training, augmentation and tool use produced distinct motor coordination signatures. Tool use strengthened existing finger coupling and progressively compressed task-space variance, consistent with refinement of an established control policy. Augmentation, in contrast, departed more strongly from the initial hand coordination space, drove broader movement exploration that extended to the elbow and shoulder, and increased the range of movements within the null-space, while task-space variance remained elevated. Crucially, this null-space variability became increasingly structured to stabilise end-effector position. Augmentation did show some adaptation-like features, with inter-finger coupling decreasing early in training and then stabilising, but these sat within a broader pattern of change not easily explained by recalibration alone. Augmentation therefore appears to engage a qualitatively different learning process: instead of tightening an existing coordination structure, participants explored the expanded motor space introduced by the device and progressively organised this exploration into coordinated movements that stabilised the effector.

### Biological–robotic interaction, not augmentation alone, determines the scope of motor restructuring

Independent augmentation removes the collaborative demand between the robotic and biological digits. Participants used the Third Thumb alone (Fig. 1C, Supplementary Vid. 3) to extract a peg from a fitted holder and insert it into another, with no biological finger contribution to the grasp (*31*, *32*). This task allowed us to test whether controlling an added effector without biological finger collaboration was sufficient to drive the upper-limb restructuring observed in the Interaction task, in which participants grasped cylindrical objects by coordinating the robotic digit with one of their biological fingers. Therefore, we repeated the analyses above, comparing the Interaction and Independent tasks directly.

Performance improved comparably across both augmentation tasks (Independent: *F*(4,80.42) = 94.99, *p* < .0001; Day 1 vs. 5: paired *t*(84) = − 17.26, *p* < .0001; Task: *F*(1,40.78) = 1.01, *p* = .320; Day × Task: *F*(4,162.05) = .89, *p* = .472; Fig. 2A), with no evidence of different learning rates (paired *t*(21) = − 1.68, *p* = .109, BF10 = .74; Fig. 2C). Wrist smoothness also increased similarly in both tasks (Day: *F*(4,154.66) = 128.56, *p* < .0001), and learning rates were anecdotally comparable (paired *t*(16) = −1.11, *p* = .284, BF_10_ = .42; Fig. 2D). Speed–accuracy slopes, however, followed different time courses in the two tasks (Day × Task: *F*(4,148.70) = 2.96, *p* = .022). On Day 1, slopes were steeper in the Independent than in the Interaction task (paired *t*(132) = 3.11, *p* = .002), but they did not differ significantly on Day 5 (paired *t*(133) = −1.23, *p* = .220). In the Independent task, faster movements were associated with greater trajectory error on Day 1 (slope against zero: *t*(50.4) = 7.64, *p* < .0001) but not on Day 5 (slope against zero: *t*(91.2) = .65, *p* = .515; Fig. 3A,C). Therefore, in both augmentation tasks an initial speed–accuracy trade-off was no longer significant by the end of training, in spite of continued performance gains. This shared pattern suggests that controlling the Third Thumb itself, whether alongside or independently of the biological fingers, involved acquiring a new control policy, which is consistent with de novo learning. Performance and smoothness were closely matched across the two augmentation tasks, although the trade-off was initially steeper when the robotic finger operated independently of the biological fingers.

The two augmentation tasks also differed in how limb coordination restructured during early training. Although the biological fingers did not contribute to the Independent grasp, using the robotic digit still engaged arm exploration (Fig. 4D). On Days 1-2, elbow and shoulder deviation from the late-training reference followed different trajectories across tasks, with greater deviation in the Independent task (Day × Task × Body part: *F*(4, 318.32) = 4.20, *p* = .002; all *p* < .05), whereas wrist deviation was greater in the Interaction task (all *p* < .05). Comparable learning trajectories therefore arose through distinct patterns of upper-limb restructuring, suggesting that even though participants controlled the same device, the presence or absence of biological-artificial finger collaboration shaped a different exploration space.

To further characterise this restructuring, we quantified the range of motion (ROM) at the wrist, elbow, and shoulder joints (Fig. 6A). ROM decreased significantly from Day 1 to 5 in the Interaction, Tool and Independent tasks, but at different rates (Day × Task: *F*(8,737.44) = 7.85, *p* < .0001). ROM was initially larger in the Independent than in the Interaction and Tool tasks (all *p* < .001), reversing by Day 4 (all *p* < .001). The slope in the Independent task was roughly 2.5 times steeper than in the Interaction (Interaction: *b* = −.21; Independent: *b* = −.54; Interaction vs. Independent: *t*(764) = 6.19, *p* < .0001) and Tool tasks (Tool: *b* = −.19; Independent vs. Tool: *t*(764) = −6.50, *p* < .0001). The Interaction and Tool slopes did not differ significantly (Interaction vs. Tool: *t*(766) = −.37, *p* = .929). The steeper reduction in the Independent task may thus reflect a more rapid convergence on a motor solution when biological–robotic finger collaboration is absent, in accordance with a more constrained solution space.

**Figure 6.**
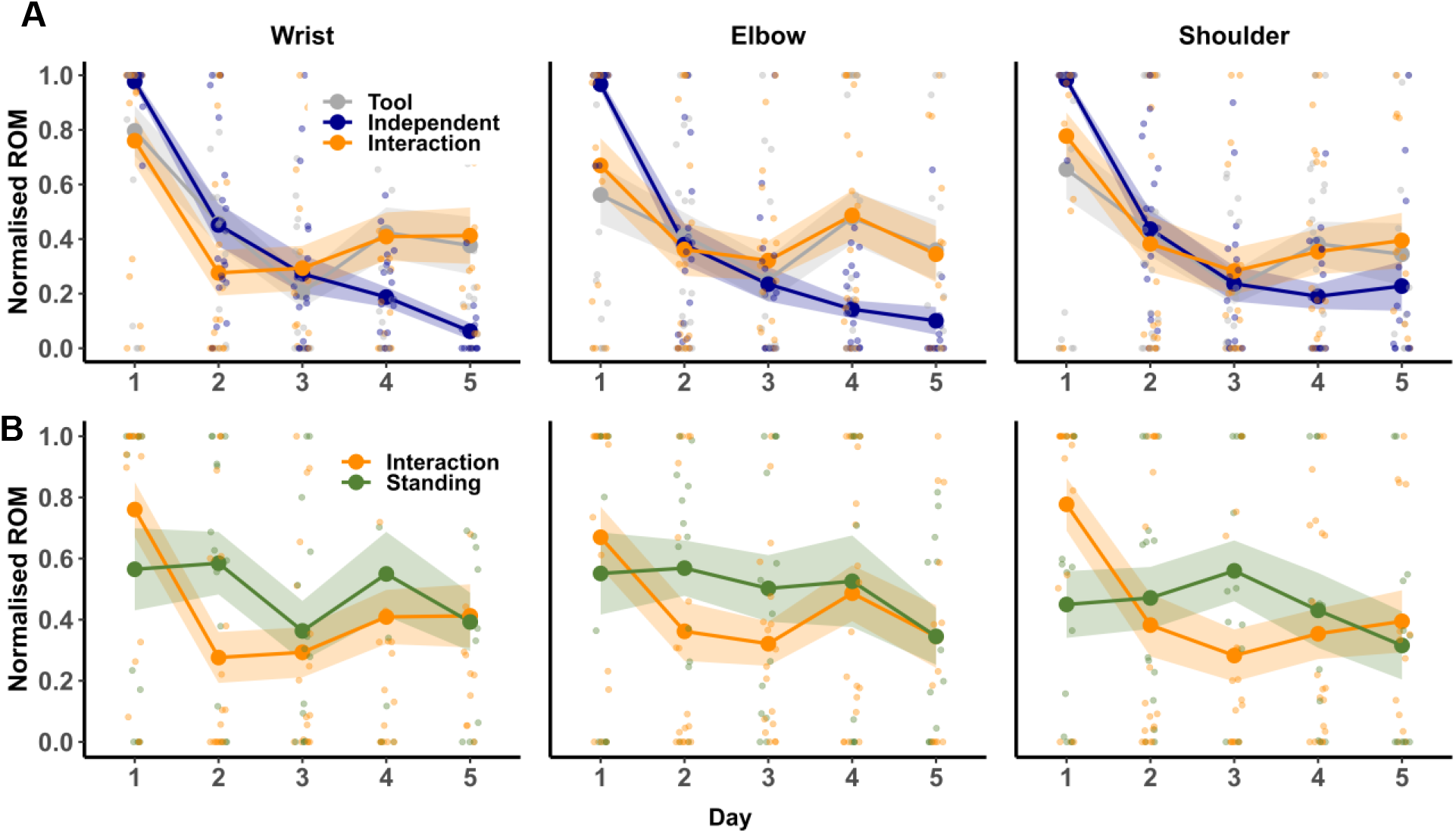
Upper-limb range of motion (ROM) across augmented training. Normalised range of motion (ROM) of the wrist (left), elbow (middle), and shoulder (right) across the five training days. **A,** Interaction, Tool, and Independent tasks, all performed seated. ROM followed similar trajectories in the Interaction and Tool tasks at all three joints and decreased more steeply in the Independent task. **B,** Interaction task performed seated and standing. Other figure annotations are as detailed above.

The narrower exploration space for the Individuation task was also reflected in the upper-limb joint variability analysis (Fig. 5B). Both task-space (Day × Task: *F*(4,151.17) = 6.06, *p* < .001; Day 1 vs. 5: paired *t*(154) = 5.52, *p* = .040) and null-space variance (Day × Task: *F*(4,145.02) = 10.77, *p* < .0001); Day 1 vs. 5: paired *t*(148) = 5.80, *p* < .0001) declined throughout training in the Independent task, with both components significantly higher than in the Interaction task on Day 1 (all *p* < .0001). The balance between these components likewise evolved differently (Fig. 5D). The task-stabilising synergy index *ΔV* declined across training in the Independent task (Day × Task: *F*(4, 146.57) = 6.94, *p* < .0001; Day 1 vs. 5: paired *t*(150) = 4.16, *p* < .0001), with significant differences between tasks at Day 1 (paired *t*(148) = 3.96, *p* < .001) and Day 5 (paired *t*(147) = −3.11, *p* = .002). The declining synergy index indicates that Independent augmentation compressed variability uniformly, without discriminating between task-relevant and task-irrelevant directions. This is more consistent with general adaptation of an existing control policy, which affects all dimensions at once, than with learning the task’s redundancy structure. By contrast, the Interaction task produced progressively stronger task-stabilising synergies, suggesting that it may be the collaborative demand between biological and robotic digits, not merely controlling an additional effector, that drives the restructuring of motor synergies. However, this result should be interpreted cautiously, as the biological thumb tip was used as the end effector in both tasks for comparability, while it did not contribute to the grasp in the Independent task (see Discussion for further details).

### Augmented performance is maintained across postural change

We next tested whether expanding the potential solution space, by extending participants’ postural range, would alter performance and motor exploration. To this end, we contrasted the Interaction task performed seated versus standing (Fig. 2A, Supplementary Vid. 5). Performance improved in both postural scenarios, with no significant difference between them (Standing: *F*(4,75) = 38.99, *p* < .0001; Day 1 vs. 5: paired *t*(60) = − 9.91, *p* < .0001; Task: *F*(1,176) = .43, *p* = .514; Day × Task: *F*(4,176) = 1.51, *p* = .202), anecdotally corroborated by their similar log-linear learning rates (paired *t*(16) = −1.54, *p* = .143, BF_10_ = .67). Wrist kinematics were likewise not significantly different across postural contexts (Fig. 2B). Despite a significant smoothness difference at Day 4 (Day × Task: (*F*(4, 126.76) = 2.61, *p* = .039; paired *t*(132) = 2.89, *p* = .005), smoothness learning rates did not differ between tasks (paired *t*(9) = .08, *p* = .936, BF_10_ = .31, Fig. 2D), suggesting that this difference reflected minor day-to-day fluctuations rather than a meaningful divergence in motor refinement across postural contexts. Speed-accuracy slopes did not differ significantly between postures across days (Day × Task: *F*(4,124.47) = 1.78, *p* = .138; Fig. 3A,C). Taken together, these results suggest that standing largely preserved the learning trajectory for augmented performance and wrist kinematics.

Although standing preserved performance and wrist kinematics, the extended postural range could still have changed how the rest of the body contributed to the task. ROM (Fig. 6B) and trunk compensation (Fig. 7A) both showed significant Day × Task interactions (ROM: *F*(4, 430) = 4.38, *p* = .002; trunk: *F*(4, 143) = 2.63, *p* = .037). On Day 1, ROM was smaller when standing than when seated (*t*(429) = 2.61, *p* = .009), with no difference in trunk compensation (*t*(132) = 1.53, *p* = .128), consistent with participants restricting arm motion when standing. By Day 3, both ROM and trunk compensation were greater when standing (ROM: *t*(428) = −2.18, *p* = .030; trunk: *t*(132) = −2.85, *p* = .005), suggesting participants drew on the extended postural range by relying more on the trunk and arm. On Day 5, neither measure differed between standing and seated (ROM: *t*(416) = .36, *p* = .718; trunk: *t*(126) = .313, *p* = .755). To test whether the greater range of motion when standing interfered with the task, we decomposed arm joint angle variance into task-related space and null-space components (Fig. 7A). Both task-related space and null-space variance were greater in Standing than in Interaction (*V_ORT_*: *F*(1, 138.65) = 5.30, *p* = .023; *V_UCM_*: *F*(1, 131.70) = 17.00, *p* < .0001). When these components were combined to assess end-effector stability (Fig. 7C), the task-stabilising synergy index *ΔV* was significantly greater in Standing than in Interaction (*F*(1, 131.07) = 15.66, *p* < .001), indicating that the increased joint angle variability was organised to preserve task performance through a disproportionate expansion of null-space variance.

**Figure 7.**
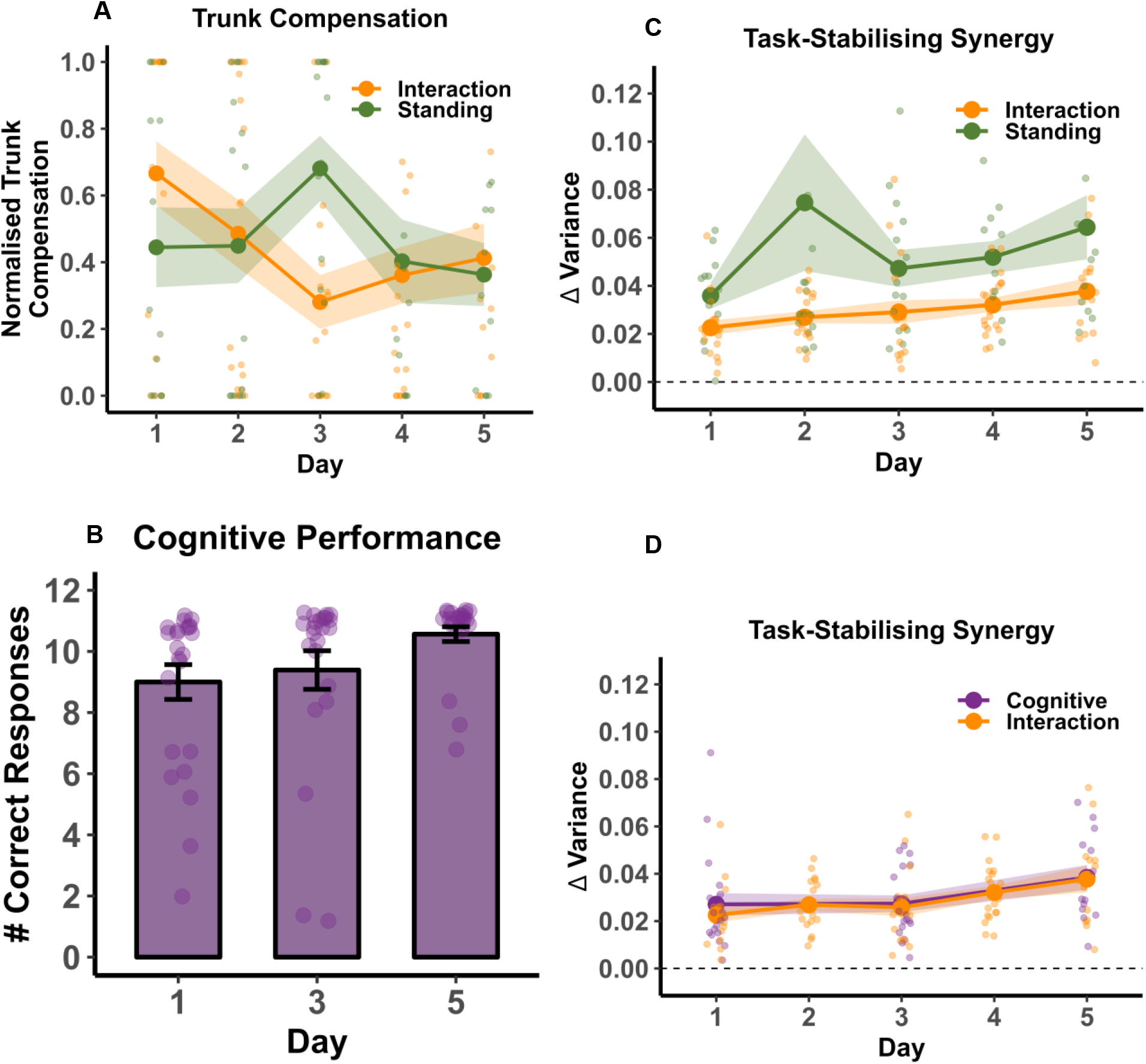
Postural, but not cognitive, demands altered task-stabilising synergy. **A**, Normalised trunk compensation in the Interaction task performed seated and standing. Trunk compensation was greater when standing on Day 3 and did not differ on Days 1 and 5. **B,** Number of correct responses in the arithmetic task increased significantly across training days. **C, D,** Task-stabilising synergy index (*ΔV* = *V*_UCM_ −*V*_ORT_) in the seated Interaction task compared with the Standing (C) and Cognitive (D) tasks. *ΔV* was greater in the Standing than the Interaction task (C), and did not differ between the Interaction and Cognitive tasks (D). Other figure annotations are as detailed above.

Together, these results suggest that expanding the solution space altered motor exploration but not performance. Participants exploited biomechanical redundancy when standing, structuring the additional joint variability to preserve end-effector stability.

### Cognitive robustness and growing embodiment signal motor repertoire integration

Cognitive load has previously been shown not to impair task performance with the Third Thumb (*23*). Nonetheless, here we asked whether it might implicitly alter how movements are executed, which would be visible in the kinematics even when performance is preserved. Participants performed the Interaction task while simultaneously performing an arithmetic task, a paradigm commonly used to assess cognitive load in prosthesis research (*23*, *35*, *43*, *44*). Task performance improved comparably across days in the Cognitive and Interaction tasks (Cognitive: *F*(2,38.96) = 36.68, *p* < .0001; Task: *F*(2,97.91) = 3.57, *p* = .062; Day × Task: *F*(2,97.91) = 1.30, *p* = .278; Supplementary Fig. 1A). Speed–accuracy slopes also did not differ significantly between tasks (Day × Task: *F*(2,70.49) = 2.61, *p* = .080, Fig. 3, Supplementary Fig. 1B,C). This result does not, however, discriminate between learning processes. De novo learning would predict dual-task interference early in training that diminishes with practice (*26*), which this design was not positioned to reliably detect.

Added cognitive load did not change how wrist smoothness, overall movement variance or task-stabilising synergies developed with practice (Figs. 2B, 7D; Supplementary Fig. 2B). Wrist smoothness increased similarly across tasks (Day: *F*(2,79.57) = 75.56, *p* < .0001; Task: *F*(1,67.24) = .38, *p* = .539; Day × Task: *F*(2,67.31) = .16, *p* = .852; Fig. 2B). Variance organisation also mirrored the Interaction task (Supplementary Fig. 2B). The task-stabilising synergy index *ΔV* increased equally in both tasks (Day: *F*(2,72.81) = 10.33, *p* < .001; Task: *F*(1,70.54) = .03, *p* = .860; Day × Task: *F*(2,70.58) = .26, *p* = .775; Fig. 7D). Together, these findings indicate that despite the more prominent de novo signatures of augmentation learning, the concurrent arithmetic task did not measurably disrupt augmented motor performance, consistent with previous reports of minimal dual-task interference during body-machine interface control (*23*, *35*, *43*, *44*).

Cognitive performance also improved across training (Friedman test: *χ²* (2) = 15.86, *p* < .001), with correct responses increasing from Day 1 to 5 (Wilcoxon signed rank test, *p* = .007; Fig. 7D). Concurrent gains on both tasks may indicate that device control became increasingly automatic with practice, freeing cognitive resources for the arithmetic task. This would be compatible with the typical course of explicitly acquired motor skills, which draw on cognitive resources early in learning and become automatised with practice (*45*, *46*), as previously observed with the Third Thumb (*30*). However, as the arithmetic task was not performed without the device at baseline, improvements in cognitive performance cannot be separated from practice effects on the arithmetic task itself, and should be taken as suggestive rather than direct evidence of increased automaticity.

Beyond behavioural and kinematic outcomes, we finally assessed whether training with the robotic finger was accompanied by shifts in subjective sense of embodiment (Fig. 8). Participants rated four dimensions (Body Ownership, Agency, Body Image, and Somatosensation) on the first and fifth day of training. Perceived embodiment increased significantly relative to control items for each of the four dimensions (Body Ownership: (*V* = 187, *p* = .002; Agency: *V* = 172.5, *p* = .002; Body Image: *V* = 192, *p* = .002; Somatosensation: *V* = 188.5, *p* = .002), indicating that motor learning with the robotic digit was paralleled by a broad increase in subjective embodiment.

**Figure 8.**
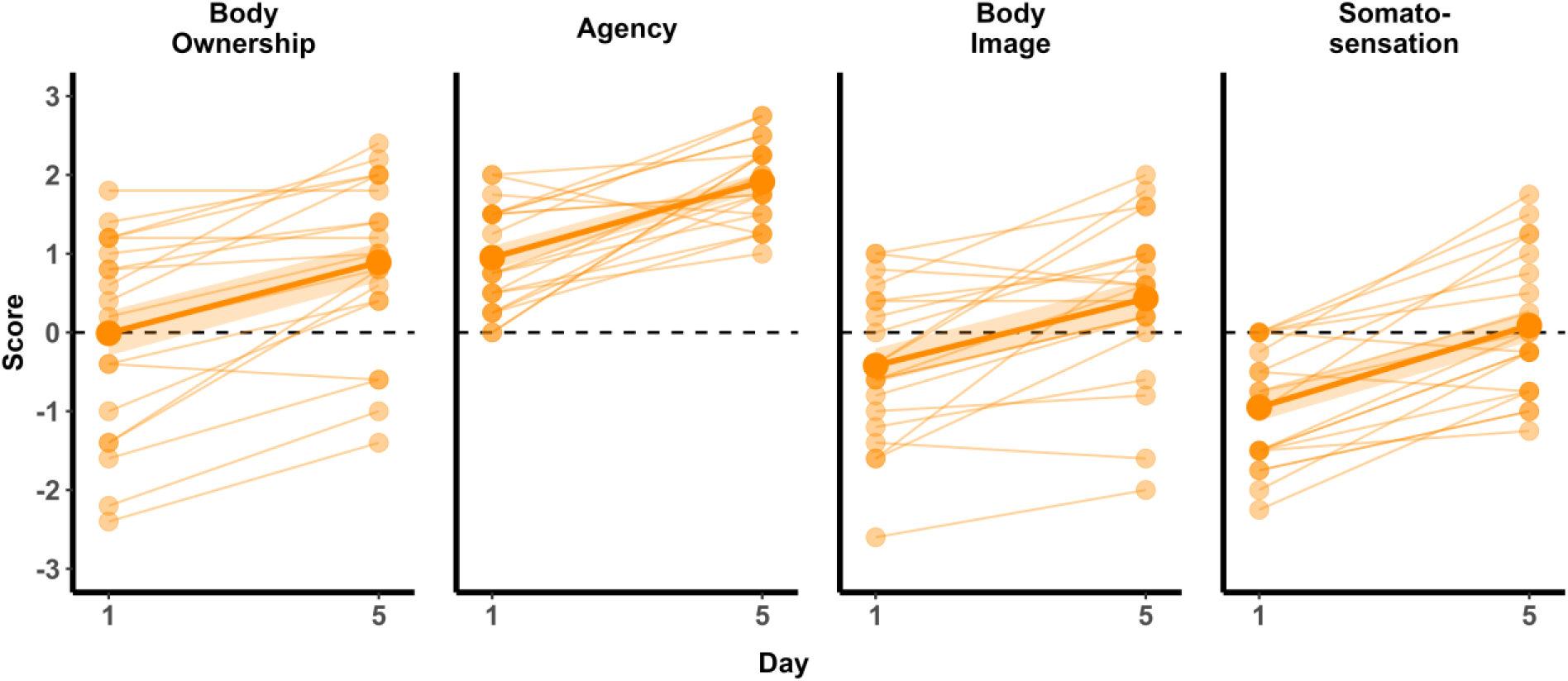
Subjective embodiment increased across training. Ratings for four embodiment dimensions (Body Ownership, Agency, Body Image, Somatosensation) across training days, on a 7-point scale from −3 to 3. Ratings increased from Day 1 to Day 5 on all four dimensions. Dots: individual participants; Coloured lines: group means, Shaded regions: SEM; Dashed line: zero.

Together, these results suggest that augmented motor control was largely maintained under concurrent cognitive demands, across performance, kinematic and synergy-level measures. Together, preserved performance under cognitive load, structured kinematic compensation under postural challenge, and increasing subjective embodiment across training are consistent with progressive integration of the robotic finger into the user’s motor repertoire.

## Discussion

Wearable robots are typically evaluated by how quickly users reach proficient performance (*1*, *2*, *47*, *48*), yet little is known about how the user’s own movements change to accommodate them. Tracking naïve users across five days of training with a wearable robotic thumb, we found that even though the device is small and attached locally to the side of the palm, users had to adjust coordination across the entire upper limb as they learnt to use it proficiently. This process of motor learning did not stabilise around an initial motor solution but continued to rearrange across days. Over training, finger coordination progressively departed from its initial structure. Task-stabilising synergies, reflecting covariation among joints that preserves end-effector position, increased across the kinematic chain from biological thumb to shoulder, while movement exploration engaged the elbow and shoulder, as well as the wrist. Previous work showed that augmentation training alters coordination among the biological fingers (*23*, *24*); our results show that these changes extend across the whole upper limb. Together, these findings suggest that proficient integration of a wearable augmentation device involves exploration and refinement of movement across the entire upper limb over time.

We propose that these gradual, limb-wide changes are best understood as reflecting two motor learning processes that simultaneously contribute to learning to control a SRL. Motor adaptation recalibrates an existing, or newly formed, control policy. Classically, adaptation has been studied as a process that restores performance after a change in the environment or body, while coordination remains largely within its existing structure (*25–28*), for example when stabilising an endpoint under a force perturbation. However, when a task demands a mapping that the existing policy cannot accommodate, adaptation is insufficient and may even work against performance (*49*, *50*). De novo learning, instead, forms a new control policy that changes how actions are selected and executed in a given context by broadening the available repertoire (*26*, *27*). For task-stabilising synergies, this would mean incorporating a new effector, or a new combination of joints, into the coordinated pattern that stabilises the task. For example, when an object is picked up between the Third Thumb and a single biological finger (*23*), the existing grasp policy, built around opposition with the biological thumb, cannot simply be recalibrated. A new policy must be built, and grasp stability would then depend on coordination between that finger and the extra thumb. Unlike adaptation, which can occur within a single session, de novo learning typically develops over days of practice (*27*, *28*). The two processes are not mutually exclusive (*27*, *28*): coordination between the extra thumb and the biological fingers must be learned de novo, and both this new policy and the existing control of the biological fingers can then be refined through adaptation to accommodate the extra digit. To distinguish these processes, we compared augmentation with two other conditions: learning to use a novel hand-held tool, and controlling the robotic digit independently of the biological fingers.

We first compared augmentation with tool-use to test whether these changes were specific to augmentation. Some features of augmentation learning were consistent with adaptation or reflected general practice effects shared with the learning process involving using a novel tool. In the Interaction task, inter-finger coupling loosened from Day 1 to Day 2 and then plateaued. This rapid change followed by stabilisation matches the time course of adaptation, consistent with existing finger control being recalibrated to accommodate the extra digit. Other features reflected general practice: task performance improved and joint range of motion declined at comparable rates in the Interaction and Tool tasks. Muscle activation patterns also remained similar to those during tool-use, suggesting reuse of existing actions, which is compatible with either process (*26*, *28*). By contrast, several features were specific to augmentation and more consistent with de novo learning. Null-space variance and task-stabilising synergies increased while task-space variance remained stable, indicating that joint variability became progressively organised into a task-stabilising pattern (*51–53*). Wrist smoothness also improved across training. Sustained gains in movement quality over days are characteristic of de novo learning, which improves how actions are executed, whereas adaptation recalibrates existing actions to restore baseline performance without refining their execution (*26*). Faster movements were associated with greater trajectory error on Day 1 but not on Day 5, while performance continued to improve throughout, suggesting learning beyond the initial speed–accuracy relationship (*40*). Finally, finger coordination departed further from its initial structure than in tool-use, and early movement exploration, which was greater in augmentation than in tool-use, declined across training, consistent with convergence on a new solution (*28*). Tool-use, in contrast, showed decreasing task-space variance with stable null-space variance, strengthened inter-finger coupling, and no change in smoothness or speed– accuracy relationship, a profile more consistent with refining existing coordination than forming a new control policy. Therefore, although task performance improved comparably in the two tasks, we propose that the underlying learning processes are distinct.

The augmentation-specific restructuring of joint variability was not a general consequence of controlling an added effector. If it were, the Independent task, in which the robotic digit acted without biological-finger collaboration, would have produced comparable changes. Independent augmentation did share several de novo-like features with the Interaction task: its speed–accuracy trade-off was steeper on Day 1 and no longer significant by Day 5, movement smoothness improved comparably, and early elbow and shoulder exploration was greater than in the Interaction task before converging by Day 5. The way participants controlled their movements across the different upper arm joints, however, followed a different pattern. Task-related and null-space variance both declined, and the task-stabilising synergy index *ΔV* decreased across training, despite comparable learning rates. The decrease in *ΔV* is unlikely to be an artefact of the UCM endpoint: although the biological thumb does not contribute to the grasp in the Independent task, its tip still tracks the hand-mounted device, and *ΔV* was initially positive and higher than in the Interaction task, indicating that variability was structured around this endpoint at the start of training. The wrist, elbow and shoulder range of motion also declined more steeply than in the other tasks, consistent with rapid convergence on a motor solution. Because the robotic digit moves independently of the biological fingers while still contacting the palm, an interaction no existing policy can serve (*26*, *28*), a new policy was likely still required. The uniform reduction in variability therefore points to adaptation-like refinement within this newly formed policy rather than restructuring of coordination (*41*, *53*). What distinguishes the Interaction task is collaboration with the biological fingers. When biological and robotic digits must jointly stabilise a shared grasp, they form a new redundant system whose null-space the motor system must learn to exploit (*2*, *16*). Without this constraint, the motor system appears to reduce variability globally rather than restructure it.

Collaboration with the biological fingers may also shape how the device is integrated neurally. Tools and prostheses become increasingly differentiated from the biological hand representation with expertise, in visual and parietal cortex (*54*, *55*). The Third Thumb, by contrast, is used with the hand rather than by it (or to replace it), and neural evidence suggests it is integrated with the hand representation instead (*24*). After a week of training centred on Third Thumb–finger collaboration, the S1 representation of the Third Thumb became more similar to that of all five biological fingers, while its relationship with the palm, through which its sensory feedback enters, remained relatively stable (*24*). Skill training with a piano keyboard, which also altered finger use, did not produce this integration (*24*). Third Thumb training also changed how the biological fingers were used and represented. Kinematically, finger co-use decreased following Third Thumb training, which was not the case in control participants (*24*). Neurally, inter-finger similarity in S1 increased in both groups, consistent with earlier Third Thumb findings (*23*), but more so after keyboard training (*24*). Keyboard training therefore changed the biological hand representation more, whereas Third Thumb training changed how the device was integrated with it, suggesting that this integration reflects how the device is used together with the biological digits rather than altered finger use alone. The restructuring of joint variability we observed in the Interaction task may be a behavioural counterpart of this process: collaboration requires the robotic and biological digits to be coordinated as a single system, which may drive both the restructuring of motor variability and the integration of Third Thumb information with the biological fingers. Subjective embodiment also increased across training, consistent with previous Third Thumb studies (*23*, *24*, *30*). Whether using a wearable device independently of the biological digits, as in the Independent task, would produce weaker cortical integration remains to be tested.

We next asked whether these changes held beyond the seated, single-task setting. The main kinematic results were maintained under altered postural and cognitive demands. When standing, participants preserved task performance and learning rate, and although joint variability increased, the additional variability was channelled disproportionately into the null-space, producing stronger task-stabilising synergies than in the seated Interaction task (*16*, *42*). Under concurrent cognitive load, task performance, smoothness gains and task-stabilising synergy trajectories did not differ significantly from the seated Interaction task, in line with previous reports that cognitive load does not impair Third Thumb performance (*23*, *31*). The Standing results also indicate that coordination varies with the context in which the device is used, with consequences for how augmentation interfaces are designed and evaluated.

Our main findings, that augmented coordination continues to restructure as the device is integrated, have three practical implications for interface design. First, calibration should be treated as a starting point rather than a fixed mapping. Null-space interfaces assume that control signals remain independent of task-relevant biological movement (*2*, *9*, *10*), yet coordination departed from its initial pattern across five days, so this independence may erode as users learn. Interfaces should therefore monitor the relationship between control signals and biological movement and update the calibration accordingly, as in co-adaptive decoder approaches (*56*). Second, controllers should accommodate exploration and multiple control policies. Users explored different joint configurations to achieve the same outcome, and such exploration supports motor learning (*42*, *57*) Interfaces could therefore constrain task outcomes rather than enforce a single early movement pattern. They should also account for whether the added effector works alongside the biological digits or independently, as these conditions produced different coordination changes. Third, evaluation should extend beyond seated, single-task performance. Because movement patterns changed when participants stood, longitudinal testing should assess coordination across use contexts alongside accuracy and task completion.

These implications are not specific to the Third Thumb. Any device that adds controllable degrees of freedom to the biological body must draw its control from a sensorimotor system that is already fully committed to moving the body (*2*). Our results suggest that the nervous system meets this demand by restructuring existing coordination over days of practice, rather than by adding a separate control channel to an unchanged system. Interfaces for supernumerary limbs, and potentially for other devices that share control with the biological body, will therefore need to be designed for a user whose coordination continues to change. Two features of our design define the scope of these conclusions. First, the clearest de novo signatures emerged when the robotic digit worked alongside the biological fingers, so our conclusions apply to interactive robotic interfaces rather than a specific device. Second, although most features distinguished augmentation from tool-use, some were shared, consistent with the Third Thumb combining properties of a tool and a body part (*58*). Testing further devices, tools and tasks will establish how far these principles extend.

This study has several limitations. First, the tasks were deliberately constrained to capture fine-grained changes in coordination, and may not capture how the device is learned and used in daily life (*30*). Second, training spanned five days, based on prior studies (*24*, *30*), and performance gains on these tasks tended to plateau by Day 4 (Fig. 2A). Whether the observed restructuring persists, or continues to develop with longer-term use and further expertise remains unknown. Third, toe control was chosen because we previously demonstrated that the Third Thumb can be used by a broad range of people (*59*), and is robust to cognitive load (*23*). Therefore, we cannot determine whether training reduced reliance on cognitive resources or whether Third Thumb control was robust to cognitive load from the outset. This distinction matters because de novo learning in discrete reaching tasks initially relies heavily on explicit strategies (*49*, *60*), so a reduction in cognitive cost across training would provide further support for a de novo account. Fourth, our results are based on a single device, tool and a limited set of tasks, hence extending them to other augmentation technologies awaits further empirical verification. Finally, we did not test for after-effects, which are expected following adaptation but not de novo learning (*27*, *60*). After-effects occur when a perturbation is removed and the learned compensation persists, producing errors in the direction opposite to those initially induced by a perturbation (*61*). Their presence would indicate remapping of an existing control policy, hence testing for them would provide a direct dissociation between the two. Therefore, our interpretation rests on the convergence of several indirect markers: time course of performance gains with practice, specificity relative to tool-use, changes in execution quality and the structure of joint coordination. These markers cannot, however, directly establish which learning process causes the restructuring of inter-joint motor coordination.

To conclude, controlling a SRL requires harvesting of latent motor capacity from a sensorimotor system, which is already committed to moving the biological limb. Over five days of training, interactive augmentation produced a restructuring of coordination across the upper limb that was not explained by general practice or by controlling an added effector alone, suggesting that the motor system accommodates a supernumerary digit by restructuring how the whole limb is controlled.

## Methods

### Participants

Twenty-six healthy participants (14 women, 11 men, 1 non-binary, m ± sd = 27 ± 5.61 years old, range 18 - 39) were recruited to take part in the study via a recruitment panel or peer referral. Due to technical issues, five participants were excluded from the kinematic analyses. Two participants completed all sessions except Day 3, and five participants completed all sessions except Day 5. EMG data were available for 20 participants. Because Day 1 recordings were missing for 3 participants in the Interaction and 4 in the Tool task (5 unique participants), the EMG analysis was restricted to the 15 participants with complete Day 1 data in both tasks. All participants were right-handed and had no previous experience with the Third Thumb. Individuals were screened for healthy BMI (range: 18.5 – 24.9), neurological conditions and medication that affects the central nervous system. To minimise learning interference, participants were instructed to refrain from practicing another motor skill, such as playing musical instruments, computer games or participating in another motor study for the duration of the experiment. All procedures were approved by the Cambridge Psychology Research Ethics Committee (PRE.2022.068) and written informed consent was provided.

### Equipment and Setup

#### The Third Thumb

The Third Thumb (Dani Clode Design) is a robotic finger designed to expand the motor repertoire of the human hand. It is worn on the ulnar side of the right palm, opposite the biological thumb (Fig. 1A, Supplementary Vid. 6). The device has two independent degrees of freedom (DoF): flexion/extension, which moves the robotic digit towards the palm, and adduction/abduction, which moves it towards the biological fingers. Each DoF is controlled by a pressure sensor placed beneath a big toe - the right toe controls flexion/extension and the left toe controls adduction/abduction. Two wrist-mounted motors actuate the digit in proportion to the applied pressure, and releasing pressure returns the digit to its resting position. Sensor thresholds were calibrated individually for each participant. No familiarisation period was provided before task performance, so skill acquisition was captured from first exposure.

#### Motion capture

Upper-body kinematics were recorded using a custom markerless motion-capture system comprising five Teledyne FLIR Blackfly S colour cameras (BFS-U3-23S3C-C; global shutter) fitted with 4-mm fixed-focal-length lenses. Images were acquired at 1.920 × 1.200 pixels and 60 Hz in the Jarvis Acquisition Tool (*62*), with an exposure time of 4 ms and 9 dB gain. The sampling rate is more than twice the highest frequency of interest, as the power of voluntary upper-limb movement lies predominantly below 10 Hz (*63*). The cameras were arranged around a calibrated capture volume of approximately 1.4 × 1.0 × 1.0 m to maximise visibility of the right upper limb and minimise occlusion (Supplementary Fig. 3). Four AprilTags positioned at the centre of the table provided spatial references for registering the task space within the reconstructed 3D volume. To ensure frame-level synchronisation between the five cameras, a 60 Hz trigger signal was sent to a master camera via an Arduino Uno, which then triggered the remaining four cameras. LEDs placed in the corner of each camera’s field of view marked behaviour onset and offset, and recordings were trimmed in TrackStudio (*64*) by detecting frame-by-frame changes in LED luminance. Cameras were calibrated after each session, separately for seated and standing conditions, using a ChArUco board (10 × 7 squares; 25-mm squares; 18.75-mm ArUco markers from the 4 × 4, 50-marker dictionary) moved through the capture volume. Intrinsic parameters and extrinsic inter-camera relationships were estimated in Anipose (*65*) using a standard (non-fisheye) pinhole camera model and refined by bundle adjustment.

Reprojection error was used to assess whether each reconstructed point was consistent with the images from which it had been estimated. For camera *c*, joint *j*, and frame *t*, the calibrated camera model *π_c_* was used to project the reconstructed three-dimensional point **X***_jt_* back into the image, and its distance from the original two-dimensional detection **u***_cjt_* was calculated as

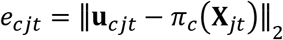

where *e_cjt_* is the reprojection error (in pixels) and ∥, ⋅.∥_2_ denotes the Euclidean norm.

Pixel errors were converted to physical distance using the estimated point depth and camera focal length, then averaged across available cameras. Errors were summarised separately for the hand, wrist, elbow, shoulders, and face. Following previously reported quality-control limits (*65*, *66*), a block was excluded if any tracked joint exceeded the following thresholds: mean reprojection error ≤ 90 mm or a maximum error ≤ 2000 mm. This ensured that subsequent analyses used only trajectories that remained geometrically consistent across views. Across retained blocks (4536/4763; 95.2%), the mean reprojection error remained below 19 mm across the training days (Supplementary Fig. 4). A detailed evaluation of tracking accuracy and quality using a similar pipeline is provided elsewhere (*64*).

#### Electromyography

Electromyographic (EMG) activity was recorded from the right forearm and upper arm during all motor tasks using a custom-made, 64-channel high-density EMG sleeve (OT Bioelettronica, Turin, Italy). Of these channels, 36 were placed over the forearm and 28 over the upper arm (Supplementary Fig. 5). Forearm flexor channels (n = 18) targeted the flexor carpi radialis, palmaris longus, flexor digitorum superficialis, and flexor carpi ulnaris. Forearm extensor channels (n = 18) targeted the extensor digitorum, extensor carpi radialis longus and brevis, and extensor carpi ulnaris. Upper-arm channels targeted the biceps brachii and brachialis (n = 12) and the long (n = 8) and lateral (n = 8) heads of the triceps brachii. A reference electrode was placed over the right olecranon. Signals were sampled at 2000 Hz via a portable Sessantaquattro+ amplifier (OT Bioelettronica, Turin, Italy) and transmitted wirelessly via Wi-Fi at 100 ms intervals to a Dell Precision 5820 Tower. Signal quality was verified before and monitored continuously throughout data collection by visual inspection of each channel’s envelope.

Channels identified as persistently noisy during quality control were excluded. Signals underwent blockwise common-mean rejection, 20–500 Hz band-pass filtering, and 45–55 Hz notch filtering. RMS and integrated EMG (IEMG) were calculated from the filtered signal. A rectified signal low-pass filtered at 2 Hz provided the activation envelope used for timing-related outcomes. For each channel and day, the MVC reference was defined as the maximum 0.5 s sliding-window RMS observed across available calibration trials. Amplitude-dependent features were normalised to this channel-specific MVC reference before analysis.

#### Questionnaires

Participants completed a 20-item Embodiment questionnaire (see *Supplementary materials Table 1*) rated on a seven-point Likert scale (1 = strongly disagree, 7 = strongly agree), comprising four dimensions: Body Ownership, Agency, Body Image, and Somatosensation (*23*, *24*, *30*) on Days 1 and 5. In addition, participants completed an Experience questionnaire with open-ended sentence-completion prompts addressing perceived utility (“I think having an extra robotic finger would help me to…”) and enjoyment of the device (“I think having an extra robotic finger would be enjoyable to…”) on the end of the study, not analysed here.

#### Experimental design

Participants completed five consecutive days of laboratory-based training with the Third Thumb, performing motor tasks adapted from previous studies (*23*, *24*, *30*, *31*). Each session began with the Tool task, followed by six Third Thumb tasks: Proportional control, Expansion, Independent, Collaboration, Interaction, and Cognitive (see *Supplementary materials*). On Days 2 and 4, the Cognitive task was replaced with the Thumb opposition task. These tasks trained both coordinated use of the device with the biological fingers and independent control of the Third Thumb. Most tasks were performed seated, however Independent, Collaboration, and Interaction were subsequently repeated while standing to assess performance generalisation across postural contexts (see *Supplementary materials Table 2*). Each task comprised five 1-min blocks, unless specified otherwise. Task order was fixed across participants and days, allowing for stable across-day comparison within task.

Participants were seated centrally at a table on a backless stool, with the table height adjusted to produce a 90° elbow angle (verified by goniometer) and the same procedure was applied for the standing configuration. Table heights were kept constant across all five days. A right-hand starting position was marked on the table and maintained throughout the study, while the left hand rested in the participant’s lap (Supplementary Vid. 7). The right arm was cleaned with alcohol, and a 64-channel EMG sleeve was fitted over the target muscle groups and secured with a skin-coloured bandage. After confirming signal quality, participants wore a white long-sleeved T-shirt over their clothing to conceal the electrodes and Third Thumb wiring, as large deviations from biologically plausible appearance adversely affected markerless motion-capture tracking. Each session began with maximal voluntary contraction (MVC) measurements followed by the Tool task, after which the Third Thumb was attached. On Day 1, toe-sensor sensitivity was individually calibrated so that the force required to fully flex or adduct the robotic finger could be comfortably maintained for the session duration. Sessions on Days 1 and 5 concluded with questionnaires.

Participants were informed that the study investigated changes in body movements and muscle activity during prolonged use of the Third Thumb. Across all tasks, the objective was to obtain as many points as possible, with the scoring system explained per task. Participants were instructed to lift and carry objects rather than drag them, whereas dropped objects could be retrieved from the table but not from the floor. All task instructions were pre-recorded to ensure standardisation. Instructions specified whether the Third Thumb should be used in isolation or in coordination with the biological fingers, however participants were free to decide which biological fingers to use. At the start of each block, participants assumed a standardised starting position: seated centrally, right hand on a marked area, left hand in their lap. Trial onset and offset were signalled by three auditory beeps.

### Data analysis

#### Kinematic reconstruction

Kinematic data was reconstructed from multi-camera recordings following procedures detailed in (*64*). Five cameras were deployed (Supplementary Fig. 3), but Camera A was excluded because it reduced calibration quality. All analyses therefore used the same four-camera array (B–E) for every session and participant. Google MediaPipe (*67–69*) was applied offline to detect two-dimensional body (33 landmarks) and hand (21 landmarks) keypoints independently in each camera view. From the body model, analyses used the right shoulder, elbow and wrist and the left shoulder. From the hand model, analyses used the 20 right-hand landmarks (four per digit), excluding the wrist, which was taken from the body model. Anipose (*65*) was used for camera calibration and combining detections across calibrated views to reconstruct three-dimensional keypoint trajectories, from which upper-limb and finger joint angles (in degrees) were derived. Before analysis, three-dimensional coordinates were transformed into a table-fixed reference frame defined by AprilTags placed on the table surface (Supplementary Fig. 3). The Third Thumb was tracked using five keypoints (right wrist, metacarpophalangeal (MCP), proximal interphalangeal (PIP) and distal interphalangeal (DIP) joints, Third Thumb tip), and its movement was reconstructed through the same multi-view pipeline. Third Thumb keypoints were detected in each camera view using a DeepLabCut model (*70*) trained on 7321 manually labelled frames, with a further 386 frames held out for testing (7707 frames in total; 95:5 split). On the held-out frames, the model achieved a localisation root-mean-square error (RMSE) of 6.41 pixels. TT kinematics are not analysed in the present manuscript.

#### Movement segmentation

Missing values in the reconstructed trajectories were filled by linear interpolation, and trajectories were then low-pass filtered with a zero-phase second-order Butterworth filter (cut-off 8 Hz) to attenuate high-frequency tracking noise. Continuous task blocks were segmented into individual action cycles using the left-right wrist trajectory along the y-axis of the task-space coordinate frame, expressed relative to the bottom right corner of the lower right AprilTag (Supplementary Fig. 3).

Peaks and troughs (local maxima and minima of the filtered trajectory) were detected with a minimum prominence of 0.6 times the standard deviation of the signal and a minimum separation of 0.3 s. A cycle was defined between adjacent troughs and was retained only if its peak exceeded an upper position threshold (170 mm) and the trajectory crossed the lower (−50 mm) and the upper threshold in sequence. Threshold crossings divided each cycle into four phases. The grasp phase extended from the downward crossing of the lower threshold at the end of the preceding cycle, through the trough, to the subsequent upward crossing of the lower threshold. Carrying extended from this crossing to the upward crossing of the upper threshold, placement from there to the peak, and return from the peak to the next downward crossing of the lower threshold. Phases shorter than 10 frames were discarded, as was the first grasp of each block, which lacked a preceding cycle. For the Independent task, grasp segments were identified as right-to-left wrist movements. Where device logs were available, grasp onset and release were refined using the left and right force-sensitive resistor (FSR) signals from the left and right Third Thumb sensors and experimenter-marked release events. Only the grasp phases were analysed, as this is the phase in which the biological fingers coordinate with the Third Thumb to secure the object, making it the most directly comparable across tasks. Cycles for which valid grasp boundaries could not be identified were excluded from further analysis.

#### Task performance and learning

Task performance was defined as the number of objects successfully transferred within a 1-minute block, logged by an experimenter via timestamped keypresses. For each participant and task, the performance was min-max normalised based on the lowest and highest performance per 1-minute block, yielding a 0–1 scale where 0 and 1 corresponded to that individual’s worst and best performance, respectively.

The participant-specific learning rate was the slope of a log-shaped curve fitted to the normalised daily block means,

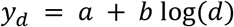

Here, d is the training day, *y_d_* is the normalised daily mean, a is the intercept, b is the learning-rate coefficient. A larger positive b indicates steeper improvement.

#### Movement smoothness, speed, and accuracy

Wrist movement smoothness was quantified with the log dimensionless jerk (LDLJ) (*36*, *37*). The three-dimensional wrist trajectory was low-pass filtered at 8 Hz using a 2^nd^ order Butterworth filter. The filter was applied bi-directionally using zero-phase forward–backward filtering to avoid phase distortion. The filtered trajectory was then differentiated. LDLJ was calculated as:

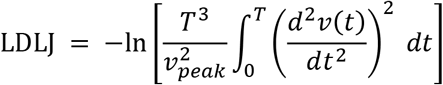

Here, T is grasp duration, t is elapsed time within the grasp, v(t) is three-dimensional wrist instant speed, and *v_peak_* is its maximum within that grasp. The second time derivative of speed was estimated by second-order finite differences, and the integral by summing squared derivative values multiplied by the sampling interval. The duration and peak-speed factors remove simple temporal and spatial scaling effects. For analysis, LDLJ values were averaged across grasps in a block and then min–max normalised within participant and task, so that 0 and 1 corresponded to the lowest and highest smoothness, respectively.

The speed–accuracy analysis paired peak planar wrist speed with trajectory error for every grasp. Peak speed was the largest frame-to-frame displacement of the filtered wrist trajectory in the x and y plane divided by the sampling interval. Peak rather than mean speed was used, following (*71*, *72*), because mean speed shares path length with the trajectory-error measure and would partly inflate their association. Trajectory error was the distance travelled beyond the shortest path:

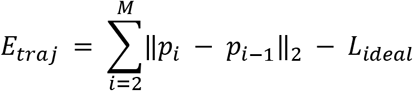

Here, *E_traj_* is trajectory error, M is the number of samples in the grasp, i indexes samples, p_i_ is the two-dimensional wrist position at sample i, ‖·‖₂ denotes Euclidean distance, and *L_ideal_* is the ideal path length. The ideal path joined the start and end positions through the wrist position at the forward–backward trough when this waypoint lay inside the grasp. Otherwise, the direct start-to-end distance was used.

Within each participant, task, and day, single-trial grasps from all available blocks were pooled and a straight line with an intercept was fitted to trajectory error as a function of non-normalised peak speed. At least three valid grasps and two distinct positive peak speeds were required. A steeper positive slope indicated a greater increase in trajectory error with increasing speed, whereas a slope near zero indicated little linear association over the observed speed range.

#### Movement pattern structure and exploration

Principal component analysis (PCA) was used to probe whether biological finger-joint coordination expressed later in training remained within the coordination space present at initial exposure. For each participant and task, a separate PCA basis was fitted to Day 1 data from 15 biological finger flexion angles: metacarpophalangeal (MCP), proximal interphalangeal (PIP), and distal interphalangeal (DIP) flexion for the index, middle, ring, and little fingers, and carpometacarpal (CMC), MCP, and interphalangeal (IP) flexion for the biological thumb. Before fitting, each angle was centred on its Day 1 mean. Angles were not scaled to unit variance and remained in degrees. Data from Days 2–5 were centred on the same Day 1 means and projected onto the Day 1 components, without refitting. The number of retained components was set as the smallest number explaining at least 85% of Day 1 variance in both tasks (k = 8; Supplementary Fig. 6). Components 9 and 10 together added only a further 3.61–6.47 percentage points. Comparable variance thresholds have been used to determine the dimensionality of finger kinematics, typically yielding 7–12 components (*73–75*).

Pairwise Pearson correlations provided a complementary measure of biological finger co-use. Within each block, grasp samples at their original sampling rate were concatenated, and correlations were calculated between MCP flexion trajectories for all ten pairs of the biological thumb, index, middle, ring, and little fingers. The MCP joint was chosen because it is tracked more reliably than distal joints by markerless pose estimation. Each pair used samples with finite values in both trajectories and required at least three such samples and non-zero variance. Correlations were Fisher z-transformed before participant-level aggregation. Greater positive correlation indicated stronger co-movement between the two fingers.

Joint deviation (JDV) quantified movement exploration relative to the participant’s own late-training solution. Each time-normalised grasp trajectory was compared with a task-specific reference obtained by averaging valid grasps from Blocks 4 and 5 on Day 5. If neither block contained a valid grasp, the last two available blocks across training were used, or the last available block if only one was available. For each block, JDV was calculated as:

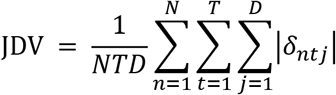

Here, N is the number of valid grasps, T is the number of normalised time points (200), D is the number of joint angles in the anatomical set, and n, t, and j index grasps, time points, and joint angles, respectively. Each grasp was time-normalised to 200 points by linear interpolation of each joint angle, providing a common temporal resolution for comparing grasps of different durations. Remaining missing values within a grasp were also linearly interpolated. A 200-point grid was chosen because some grasps contained more than 100 samples, so a coarser grid would have downsampled them. *δ_ntj_* is the angular difference, in degrees, between the observed angle and the reference angle at the same normalised time point. Periodic angles used a circular-mean reference and the shortest signed angular difference. Larger JDV therefore indicated greater departure from the late-training reference.

#### Task synergy

Uncontrolled manifold (UCM) analysis tested whether variation across a biological joint set was organised to preserve the task-relevant endpoint trajectory (*13*, *52*, *76*). The joint set comprised seven degrees of freedom: elbow flexion, shoulder elevation, shoulder-plane angle, a shoulder axial-rotation proxy, wrist flexion–extension, and two biological thumb CMC proxy angles (flexion–extension and abduction– adduction). For each grasp, endpoint motion was expressed in three trajectory-relative coordinates: distance progressed along the ideal planar path, signed perpendicular deviation from that path, and vertical position. The planar path connected the segment endpoints through the position at the wrist trough when an internal waypoint was available. The local Jacobian **J***_t_* describes how small changes in joint angles map onto changes in task coordinates at normalised time point *t*. It was estimated by least squares from frame-to-frame increments in task coordinates (*Δ***x***_k_*) and joint angles (*Δ***q***_k_*) within a local time window W*_t_*:

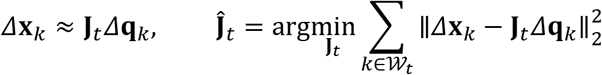

where *k* indexes increments within the window. When a window contained too few increments to determine **J***_t_* uniquely, a ridge penalty (*λ* = [X]) was added. At each normalised time point, joint-angle deviations were calculated relative to the mean joint configuration across grasps at that time point.

At each normalised time point, each grasp-specific Jacobian was decomposed by singular value decomposition, and its numerical rank was used to construct an orthogonal projector onto its joint-space task subspace. These projectors were averaged across grasps. The dimension of the common task subspace, r, was set to the median Jacobian rank across grasps (r ≤ 3, the number of task coordinates), and the leading r eigenvectors of the mean projector formed an orthonormal basis for this subspace, **V***_r_*(*t*). The UCM was defined as its (D − r)-dimensional orthogonal complement, giving the task-space and UCM projectors:

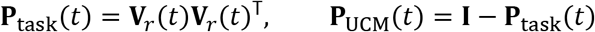

where **I** is the *D* × *D* identity matrix. Joint-angle deviations from the across-grasp mean configuration at each time point were then projected into these two orthogonal subspaces:

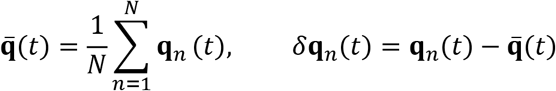

Summed squared projections were normalised by the sample degrees of freedom (*n* − 1) and the corresponding subspace dimension, yielding variance per degree of freedom in the UCM (*V*_UCM_) and in the task space (*V*_ORT_):

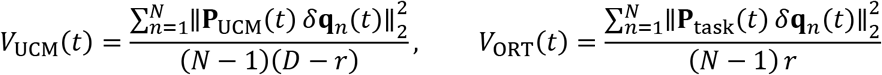

Both variance profiles were averaged across normalised time, and their balance was expressed as the synergy index *ΔV*:

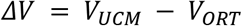

Here, *V_UCM_* is variance in joint combinations that preserve the locally modelled task coordinates, and *V_ORT_* is variance in combinations that change them. Positive values of *ΔV* indicate greater variance per dimension in task-preserving combinations, consistent with a task-stabilising synergy.

#### Range of motion

Changes in range of motion (ROM) were quantified from marker trajectories during grasp. For each filtered three-dimensional marker trajectory, the range along each spatial axis was calculated and combined as the diagonal of the three-dimensional span:

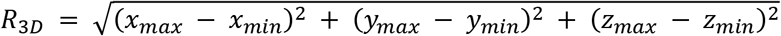

Here, *R*_3*D*_ is positional ROM in millimetres, and *x_max_*, *x_min_*, *y_max_*, *y_min_*, *z_max_*, and *z_min_* are the largest and smallest coordinates along each axis within a grasp. This quantity captures the overall spatial extent of marker movement. Trunk compensation was quantified with the same measure, applied to the midpoint between the left and right shoulder markers in the table-centred coordinate frame. Because no pelvis or sternum markers were available, this measure approximates translational trunk movement. For analysis, ROM values were min–max normalised so that 0 and 1 corresponded to the lowest and highest values, respectively.

#### EMG analysis

Electromyography (EMG) analyses were restricted to the grasp phase, using the same kinematic boundaries as the movement analyses. Kinematic start and end frames were mapped proportionally from the 60 Hz motion recordings to the 2,000 Hz EMG recordings. For each grasp and channel, root-mean-square (RMS) amplitude was calculated as the square root of the mean squared filtered EMG signal and normalised to the channel-specific MVC reference. This provided a measure of activation amplitude over the grasp.

For spatial analyses, each participant’s 64-channel MVC-normalised RMS values were retained as a recruitment map spanning the forearm flexors, forearm extensors, biceps brachii, and the lateral and long heads of triceps brachii. Within-participant recruitment maps were compared between the Interaction and Tool tasks on each day using cosine similarity: the dot product of the two channel vectors divided by the product of their Euclidean lengths. Values closer to one indicated more similar relative spatial recruitment, independently of an overall multiplicative difference in activation amplitude.

#### Subjective embodiment

Subjective embodiment was quantified from the Day 1 and Day 5 questionnaire responses in four dimensions: Body Ownership, Agency, Body Image, and Somatosensation (Supplementary materials Table 1). Items Q7, Q11, and Q15 were reverse-scored by subtracting the original 1–7 response from eight. Items were averaged within each dimension and participant, and four was subtracted to centre scores on the neutral midpoint, yielding a range from −3 to +3. A dimension score was treated as missing if any constituent item was missing. Higher scores indicated greater perceived embodiment in that dimension.

### Statistical analysis

Statistical analyses were performed in R (version 4.3.1; R Core Team, 2023). Linear mixed models were fitted with lme4 (version 1.1.35.1; (*77*)), and contrasts were computed on estimated marginal means with emmeans (version 1.8.9; (*78*)). Outliers exceeding the mean ± 3 SD were removed, and missing participant-days were left as missing rather than imputed.

Repeated measures analyses used full-factorial linear mixed models with a random effect for Participant. To compare tasks directly, combined models included Day, combined models included Day (1, 2, 3, 4, 5), Task, and, where applicable, Body part (wrist, elbow, shoulder), together with their interactions, as fixed effects. Planned comparisons (Day 1 vs Day 5 within each task, and between tasks on Day 1 and 5) were specified a priori and tested with t-tests on estimated marginal means without correction for multiple comparisons. All other post hoc pairwise comparisons were Tukey-corrected. Differences in learning rates between tasks were assessed with paired t-tests.

Speed–accuracy slopes were analysed with a linear mixed model including Day (1, 2, 3, 4, 5) and Task as fixed effects, with a random slope and a random intercept for Day and Participant, respectively. To determine whether trajectory error increased significantly with peak wrist speed, estimated marginal means for each task and day were tested against zero, with false discovery rate (FDR) correction for multiple comparisons.

For null results of interest, Bayes factors (BF₁₀) were computed using a Cauchy prior (width *r* = 0.707), with BF₁₀ < 1/3 interpreted as support for the null, BF₁₀ > 3 as support for the alternative, and intermediate values considered inconclusive (*79*).

For the embodiment questionnaire, responses were averaged within each category, resulting in a single composite score per category. Change scores (Day 5 − Day 1) were then computed for each participant separately for each of the four embodiment categories (Body Ownership, Agency, Body Image, Somatosensation) and for the control items. Each category’s change score was compared with the control change score using Wilcoxon signed-rank tests, FDR-corrected across the four comparisons.

## Acknowledgements

We thank all participants for their time and commitment across the five training sessions. We thank Natalia Torres-Consul and Genji Kawakita for help with camera synchronisation, Jonathan Michaels and Andrew Pruszynski for advice on camera synchronisation, and Grace Hu, Ellie Carre, Andrew Dott, and Angus Gray for help with data collection. We thank Lucy Dowdall for helpful discussion. We thank Gary Chandler and Mark Townsend for technical support and Russel Thompson and Howard Gyton for IT support. This study was funded by UKRI’s Frontier Research Guarantee (EP/X040372/1); the Medical Research Council (MC_uu_00030/10); a studentship from EPSRC CDT in Connected Electronic & Photonic Systems, to JR; a postdoctoral grant from the Fonds de la Recherche Scientifique (FNRS) [40032870]; the Department of Psychology, University of Michigan, to RP; the European Union – NextGenerationEU, under the Italian National Recovery and Resilience Plan (PNRR), through the THE – Tuscany Health Ecosystem project, Spoke 8 (ECS00000017); the European Union – NextGenerationEU, PNRR M4C2, through the BRIEF – Biorobotics Research and Innovation Engineering Facilities project (IR0000036; CUP J13C22000400007); and the Walter Benjamin Fellowship of the German Research Foundation (DFG), project number: 545902796, to CF; European Research Council, ERC-2020-StG-949660, to JAG.

## Author contributions

Conceptualization: TM, EJ; Methodology: EJ, JR, CF, VP, PK, GE; Investigation: EJ, JR, VP, ZM, PK, MG, RP, ELR, GE; Visualization: EJ, PK; Funding acquisition: TM; Project administration: TM, EJ, PK; Supervision: TM, EJ; Writing – original draft: EJ, PK; Writing – review & editing: EJ, PK, TM, JAG, HD.

## Data, code and materials availability

The code is available via GitHub at https://github.com/EmaJugovic/nullspace. The data and materials supporting the findings of this study are available within the article, in the Supplementary Information, or via GitHub.

## Supplementary Information

### Supplementary Figures

**Supplementary Figure 1.**
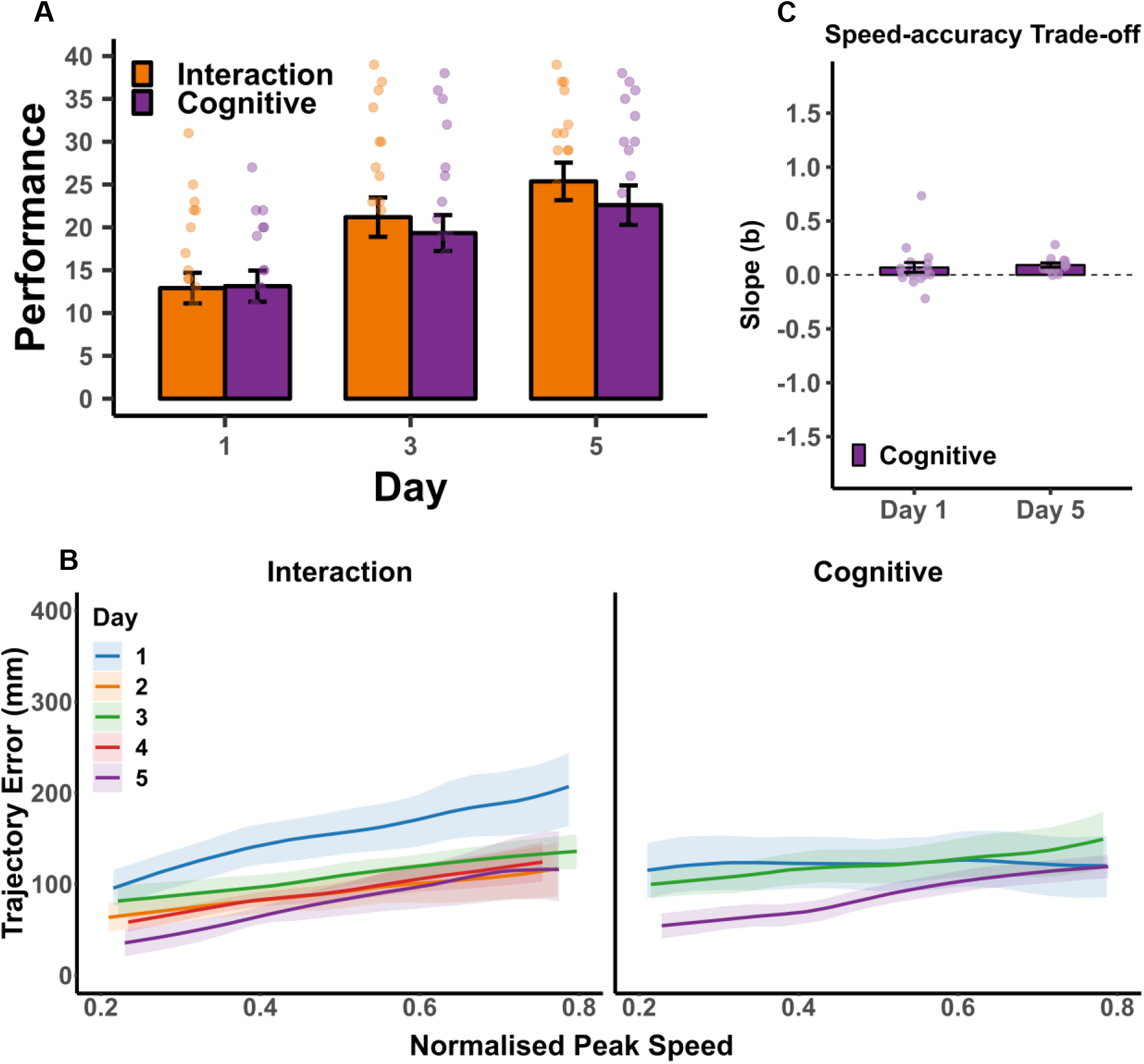
Performance and speed–accuracy trade-off in the Interaction and Cognitive tasks. **A,** Raw performance (number of objects successfully transferred) across five days of training. **B,** Trajectory error as a function of peak wrist speed on Days 1 and 5. **C,** Speed–accuracy slopes (b) in the Cognitive task on Days 1 and 5. All figure annotations are as detailed in Figure 3.

**Supplementary Figure 2.**
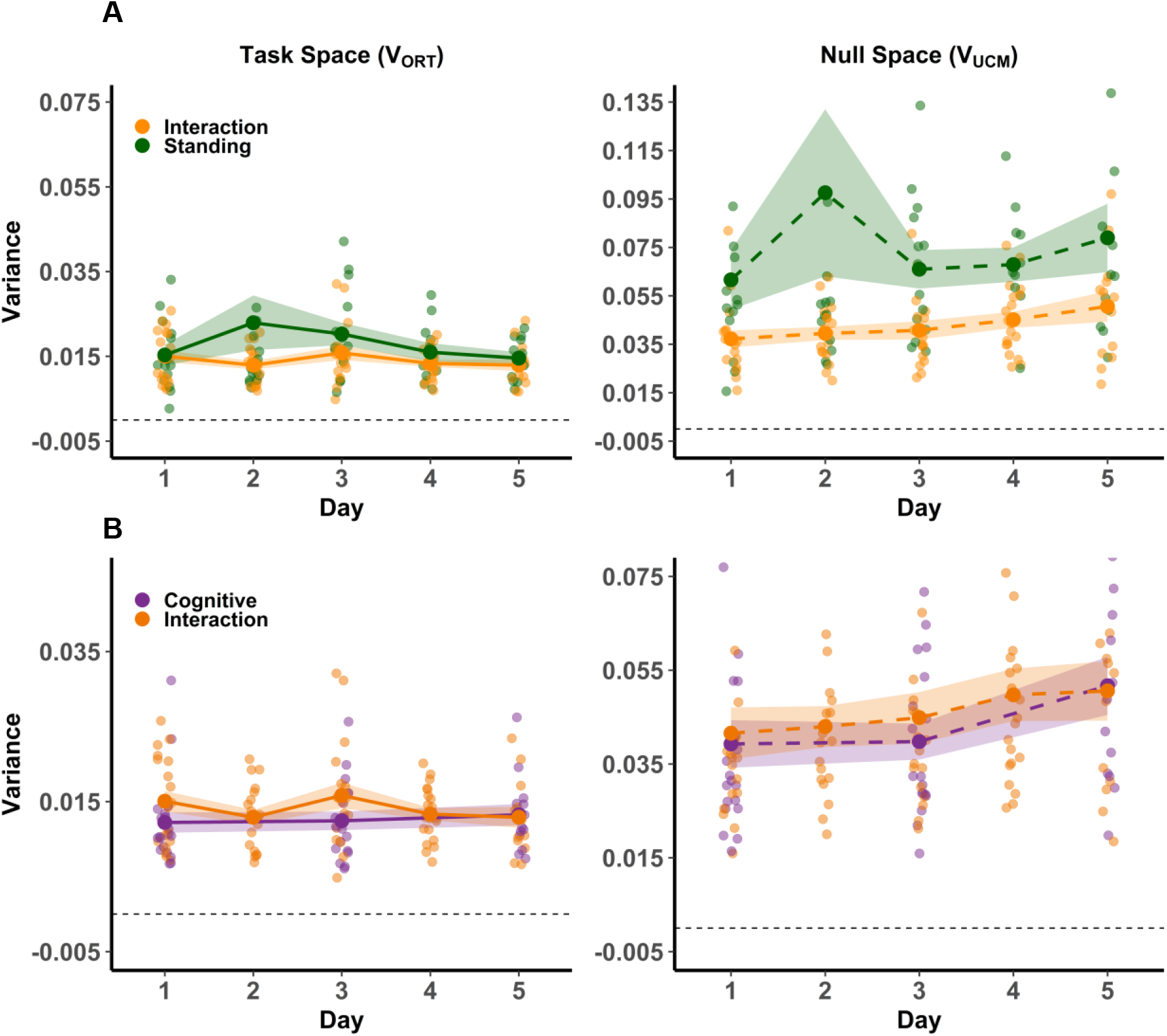
Task-space and null-space variance across postural and cognitive task contexts. Joint-angle variance was decomposed into task-space (V_ORT_, left) and null-space (V_UCM_, right) components across training days. Each component is compared between the Interaction task and the Standing task (**A**), and between the Interaction task and the Cognitive task (**B**). All figure annotations are as detailed in Figure 5.

**Supplementary Figure 3.**
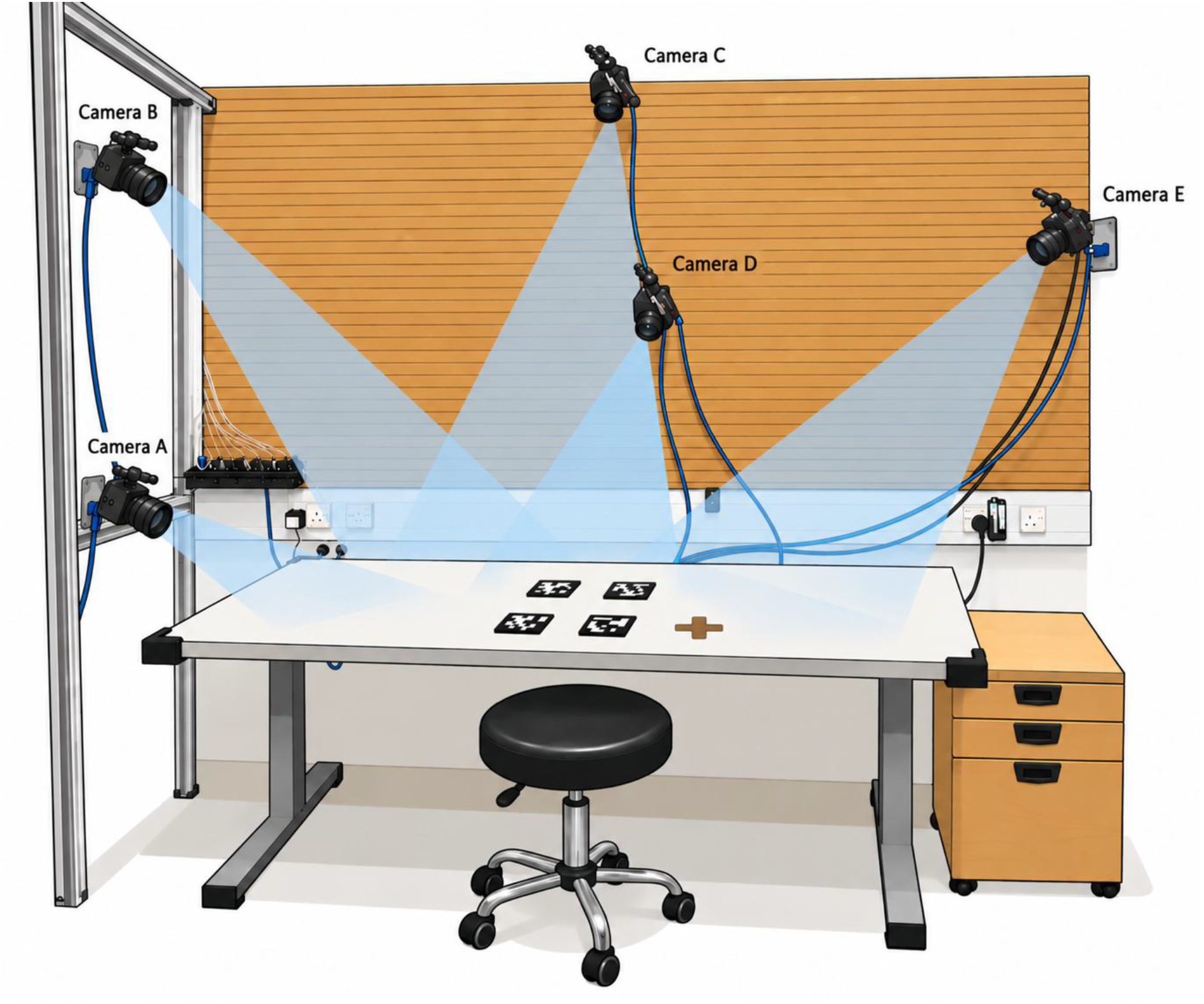
Motion capture setup for the seated condition. Five FLIR Blackfly S cameras (A–E) were positioned around the workspace to capture right hand and upper-limb kinematics. Cameras A and B were mounted on an aluminium frame to the left of the participant, cameras C and D were mounted above and behind the workspace, camera E was positioned to the right. Blue shading indicates approximate camera fields of view. Four AprilTag workspace reference markers and the participant’s starting position (cross) are visible on the table surface. The participant sat on a height-adjustable stool facing the workspace. Camera A was excluded from the final kinematic reconstruction, reducing the analysed array to four cameras (B–E).

**Supplementary Figure 4.**
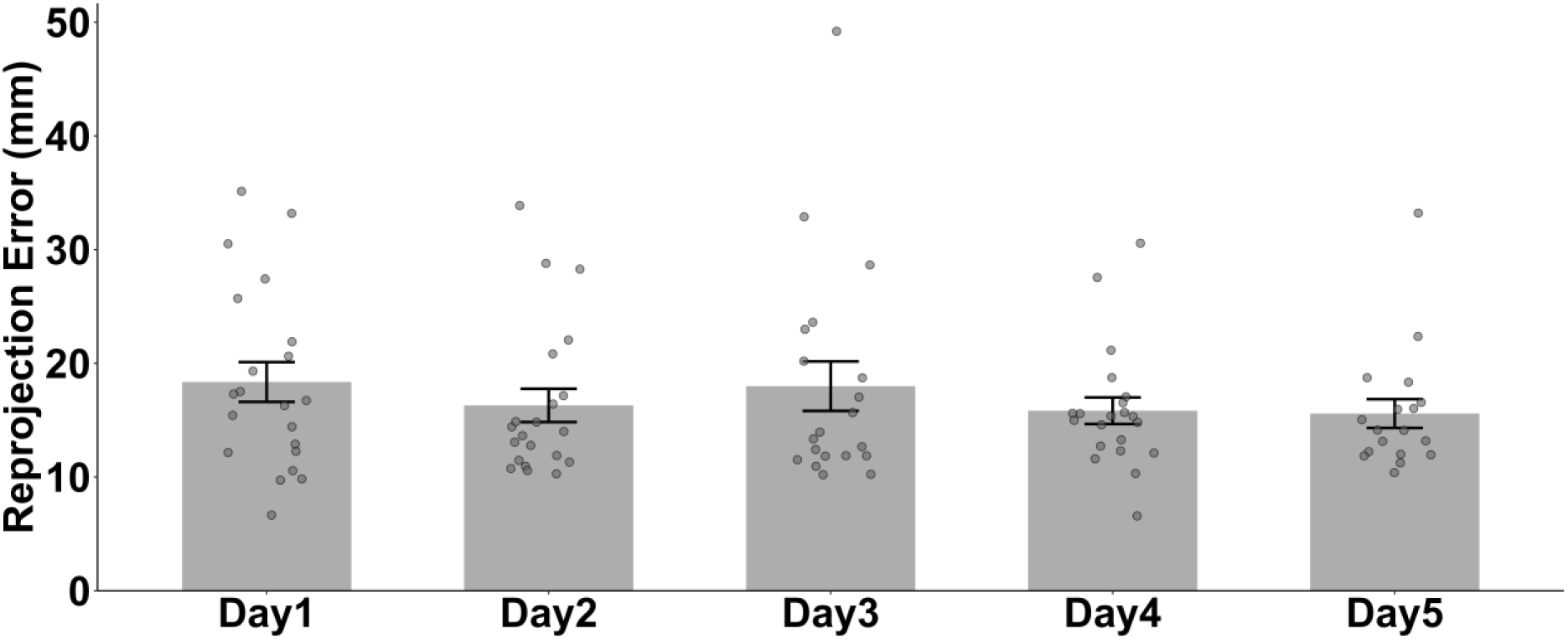
Reprojection error across training days. Mean reprojection error of the triangulated keypoints for each training day, after quality control. Pixel errors were converted to millimetres using each keypoint’s estimated depth and the camera focal length, and averaged across cameras. For each participant, errors were averaged across joints, blocks and tasks, giving one value per participant per day. Only trials that passed the quality-control criteria (mean error ≤ 90 mm and maximum error ≤ 2000 mm for every tracked joint) are included. Dots: individual participants; Lines: group means; Shaded regions: SEM.

**Supplementary Figure 5.**
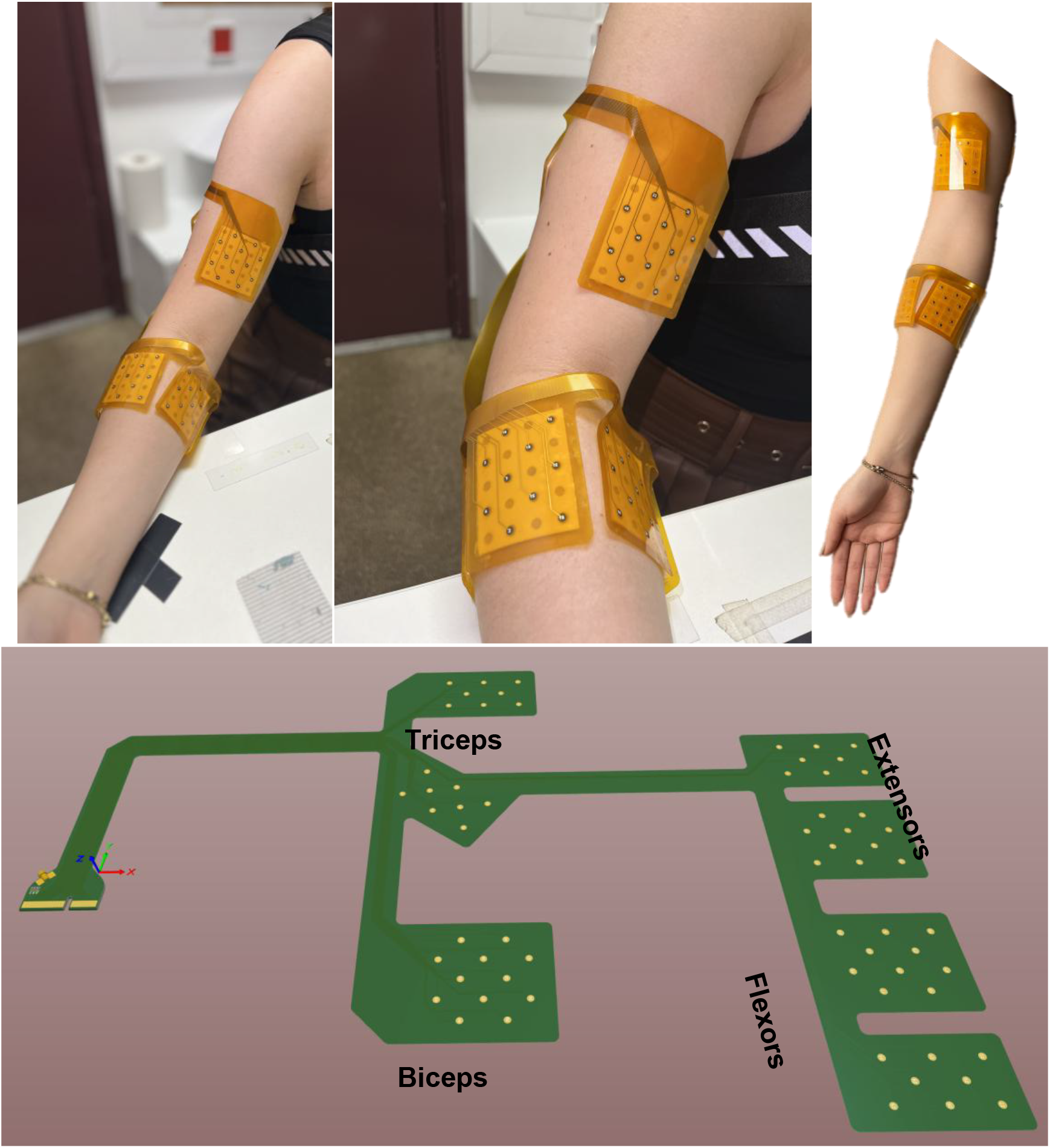
EMG electrode array design and placement. **A**, 64-channel surface EMG array on a participant’s right arm, seated at the task table. **B,** Close-up of the upper-arm (biceps) and forearm (flexors) sections of the array. Triceps and extensors are not visible. **C,** Schematic of array placement on the upper limb. **D,** Layout of the printed circuit, showing electrode pads over the biceps, triceps, forearm extensors and forearm flexors. All pads connect to a single connector (left).

**Supplementary Fig. 6.**
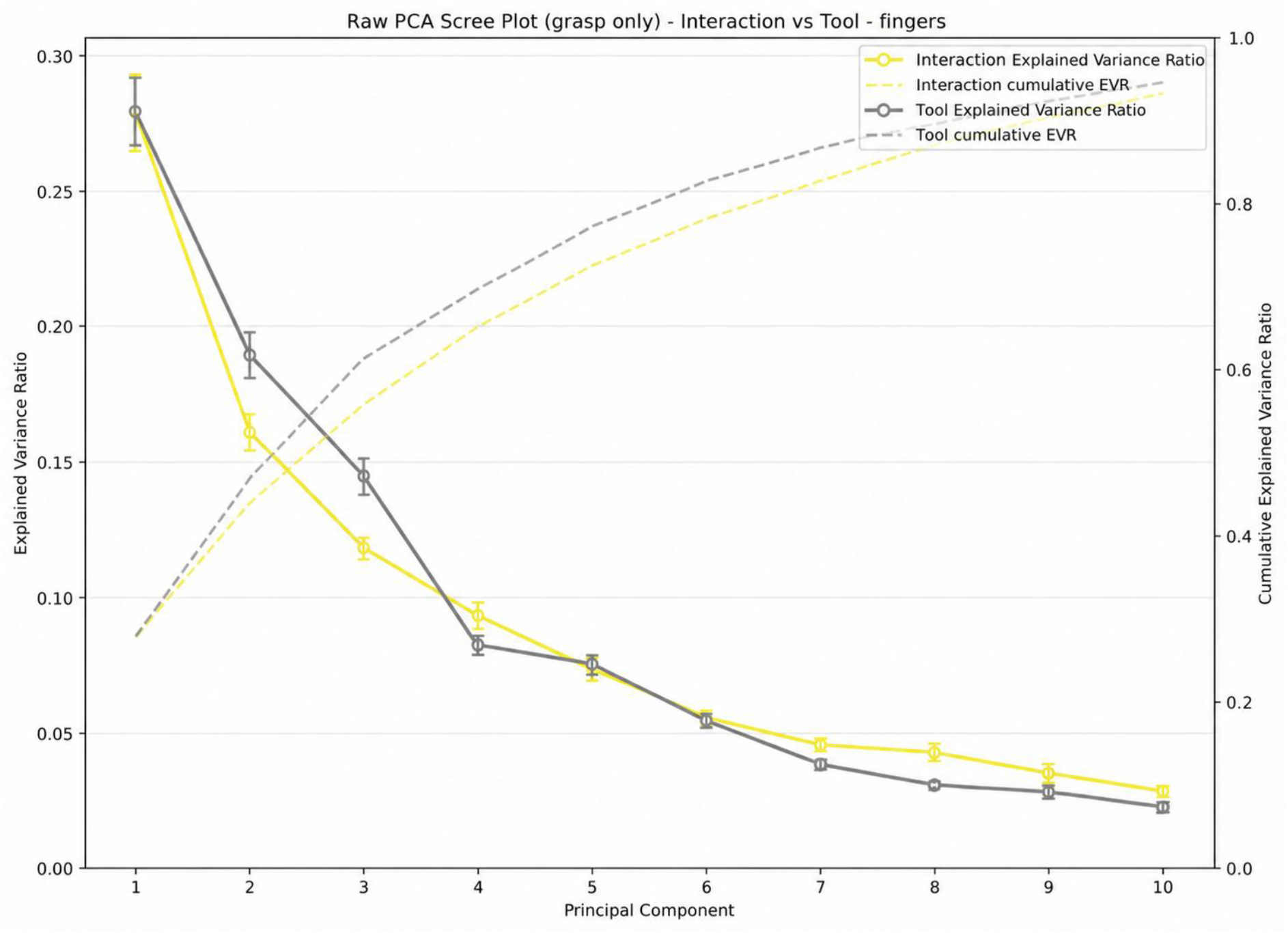
Variance explained by the Day 1 finger-coordination PCA. Principal component analysis was fitted separately for each participant and task to Day 1 grasp-phase data from 15 biological finger flexion angles (centred, unscaled, in degrees). Solid lines and circles show the proportion of variance explained by each of the first ten components (left axis), and dashed lines show the cumulative proportion of variance explained (right axis), for the Interaction and Tool tasks. Values were averaged across participants. Error bars show ±SEM. The first eight components, which were retained for further analyses, explained at least 85% of the variance in both tasks.

## Supplementary Information

### Subjective embodiment questionnaire

**Supplementary Table 1.** Embodiment questionnaire items reflecting the four embodiment dimensions.

| **Body ownership** |
|---|
| Q1: It seems like I have 3 big toes. <sup>#</sup> |
| Q2: It seems like the robotic finger belongs to me. |
| Q3: It seems like I have 6 fingers. |
| Q4: It seems I am looking at a tool fusioned to my body. |
| Q5: It seems like the robotic finger is fused with my body. |
| Q6: It seems like I have 2 heads. <sup>#</sup> |
| Q7: It seems like the robotic finger is a foreign body.<sup>*</sup> |
| Q8: It seems like the robotic finger is a part of my body. |
| **Agency** |
| Q9: It seems like I am in control of the robotic finger. |
| Q10: It seems like I can move the robotic finger at my own will, without having to pay much attention. |
| Q11: It seems like the robotic finger is moving on its own.<sup>*</sup> |
| Q12: It seems like I was causing the movement of the robotic finger. |
| **Body image** |
| Q13: It seems like I am looking directly at my own finger, rather than a prosthesis. |
| Q14: It seems like the robotic finger begins to resemble my other fingers. |
| Q15: It seems I'm wearing a tool controlled by my toes, rather than a robotic finger.<sup>*</sup> |
| **Somatosensation** |
| Q16: It seems like I can distinguish between different textures with the robotic finger. |
| Q17: It seems like I can feel temperature in the robotic finger. |
| Q18: It seems I can feel the position in space of the robotic finger. |
| Q19: It seems like I can feel the touch of an object in the location where I could see the robotic finger was being touched during object interactions/touching the object. |

<sup>\*</sup> Negative statements, <sup>#</sup> = control statements

### Experimental tasks

#### Interaction

The Interaction task assessed participants’ ability to coordinate the robotic Third Thumb with their biological fingers. Participants grasped each adhesive strip tape using the Third Thumb together with one biological finger of their choice and transferred it onto a wooden pole. They were instructed to transfer as many strips as possible within one minute. Each successfully transferred strip scored one point. The numbers of strips successfully placed and dropped were recorded as outcome measures.

#### Tool

The control Tool task mirrored the demands of the Interaction task but replaced the Third Thumb with a hand-held tool, preserving the requirement to coordinate biological fingers with an external end-effector. Participants placed as many coins as possible into designated placeholders within one minute while holding a crocheting needle between their biological fingers as they saw fit. To pick up each coin, participants used both the needle and one biological finger, grasping the coin by its rim rather than its flat surface. Outcome measures included the number of coins successfully placed and the number dropped.

#### Independent

This task encouraged participants to control the Third Thumb independently of their biological fingers. Using only the Third Thumb, participants transferred four three-dimensional-printed cylindrical inserts from one side of a table to designated holders on the opposite side. They were instructed to transfer as many inserts as possible within one minute while minimising errors. Participants followed a predefined zigzag sequence, beginning with peg 1 and proceeding numerically through the remaining three pegs. After transferring all pegs to the opposite side, they reversed the sequence to return the pegs to their original positions. This alternating sequence continued until the one-minute trial ended. The numbers of pegs successfully transferred and dropped were recorded as outcome measures.

#### Cognitive

The Cognitive task assessed augmentation performance under increased cognitive load. Its motor component was identical to that of the Interaction task: participants transferred pieces of adhesive strip tape onto wooden poles using the Third Thumb together with one biological finger. At the same time, they performed an auditory arithmetic task. Before completing the combined task, each participant underwent an arithmetic-only calibration procedure. At the beginning of each trial, participants were given a starting number, such as 127. A medium-pitched tone followed by a higher-pitched tone instructed them to add the individually selected interval to the current number. For example, with an interval of 7, 127 + 7 = 134. A medium-pitched tone followed by a lower-pitched tone instructed them to subtract the interval. Two consecutive medium-pitched tones indicated that no calculation was required. Participants reported each result aloud, and the experimenter recorded the accuracy of their responses. Each trial lasted one minute and included 11 arithmetic prompts. Trial-specific starting numbers were held constant across participants to ensure comparability. Motor performance was quantified using the outcome measures defined for the Interaction task, while arithmetic performance was quantified by response accuracy.

In addition to the core tasks, participants completed four supplementary tasks. Each task targeted a distinct aspect of Third Thumb control:

#### Proportional Control

The Proportional Control task trained participants to produce precise, graded control signals by learning the sensitivity of the Third Thumb’s control sensors. Participants transferred a weighted spherical object from one side of a table to the other as many times as possible within one minute. They grasped and transported the object using the Third Thumb together with one biological finger. The object was designed to approximate the difficulty of handling an egg without breaking it. Each successful transfer scored one point. Compressing the object sufficiently to trigger an audible click resulted in a deduction of 0.5 points, whereas dropping it resulted in a deduction of one point. This scoring scheme imposed a greater penalty for dropping than for excessive compression. The numbers of successful transfers, drops, and compression events were recorded as outcome measures.

#### Expansion

This task trained participants to expand the functional capacity of the hand using the Third Thumb. Participants grasped as many tennis balls as possible simultaneously with their biological hand and the Third Thumb, and then transferred the balls into a tray. Each trial lasted 30 seconds and permitted a single transfer. The number of balls successfully transferred was recorded as the outcome measure.

#### Collaboration

The Collaboration task trained coordinated use of the Third Thumb and biological fingers. Participants simultaneously grasped two wooden Jenga blocks. One block was grasped using the Third Thumb together with one biological finger, whereas the other was grasped using two biological fingers. Participants placed the blocks on a cross marked on the left side of the table and constructed as many 2 × 2 Jenga layers as possible within one minute. Each successive layer was oriented perpendicular to the preceding layer. Each completed layer scored 1 point, and each half-completed layer (i.e., only one block placed) scored 0.5 points.

#### Thumb Opposition

This task trained precise interactions between the Third Thumb and each biological digit. Participants sequentially performed tip-to-tip opposition between the Third Thumb and the biological thumb, index finger, middle finger, ring finger, and little finger. After completing this ascending sequence, they reversed the order, moving from the little finger back to the biological thumb. Participants alternated continuously between the ascending and descending sequences for one minute. The total number of successful tip-to-tip contacts was recorded as the primary outcome measure.

**Table 2.**
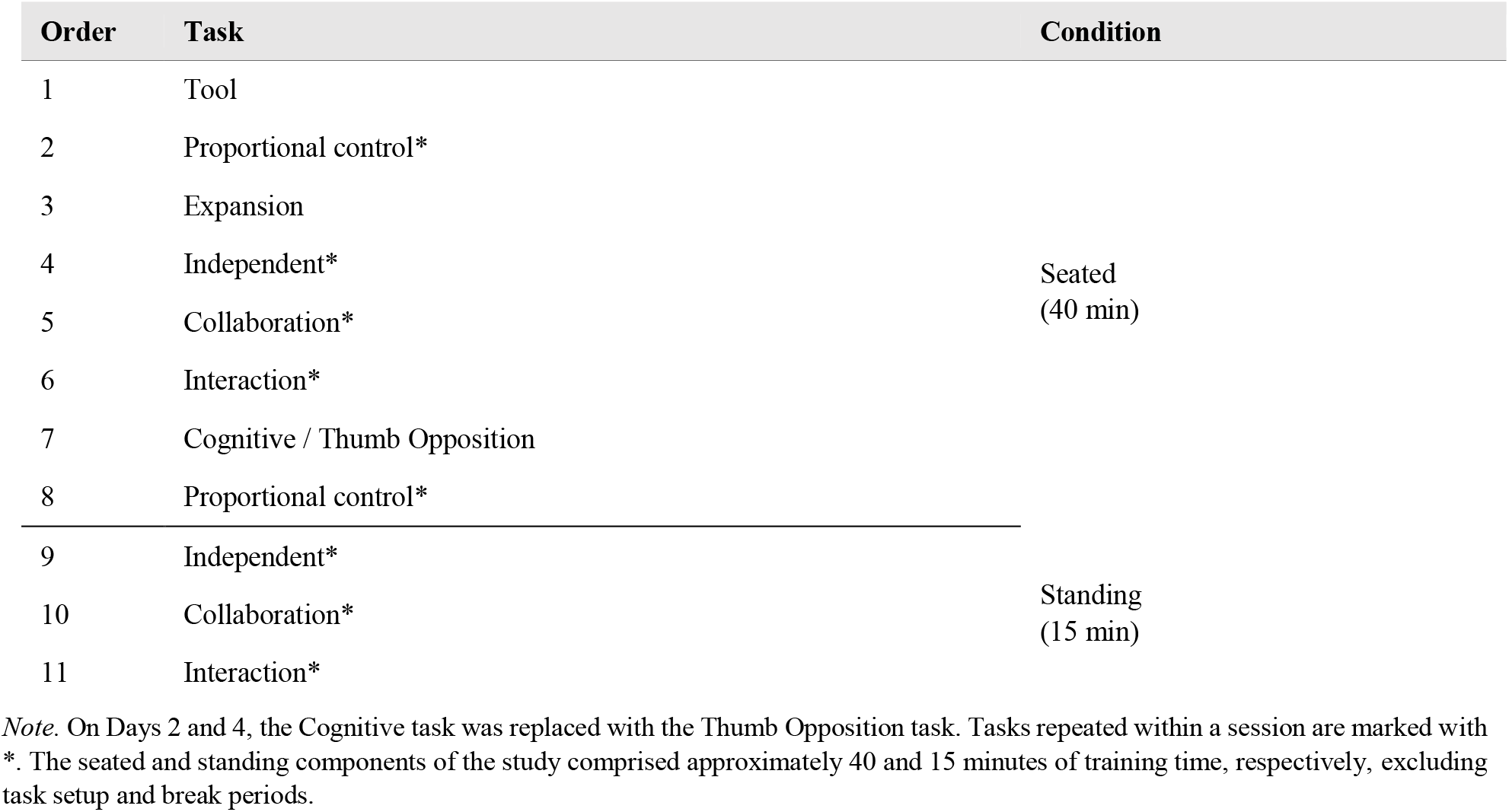
Study design and task order.

| **Order** | **Task** | **Condition** |
|---|---|---|
| 1 | Tool | Seated<br>(40 min) |
| 2 | Proportional control* |  |
| 3 | Expansion |  |
| 4 | Independent* |  |
| 5 | Collaboration* |  |
| 6 | Interaction* |  |
| 7 | Cognitive / Thumb Opposition |  |
| 8 | Proportional control* |  |
| 9 | Independent* | Standing<br>(15 min) |
| 10 | Collaboration* |  |
| 11 | Interaction* |  |

*Note*. On Days 2 and 4, the Cognitive task was replaced with the Thumb Opposition task. Tasks repeated within a session are marked with \*. The seated and standing components of the study comprised approximately 40 and 15 minutes of training time, respectively, excluding task setup and break periods.

